# Seasonal changes of airborne bacterial communities over Tokyo and influence of local meteorology

**DOI:** 10.1101/542001

**Authors:** Jun Uetake, Yutaka Tobo, Yasushi Uji, Thomas C. J. Hill, Paul J. DeMott, Sonia M. Kreidenweis, Ryohei Misumi

## Abstract

Recent progress in Next Generation Sequencing allows us to explore the diversity of airborne microorganisms across time and space. However, few studies have used consecutive short-period samples to explore correlations between the seasonal variation of the microbiota and meteorology. In order to understand airborne bacterial community dynamics over Tokyo, including fine-scale correlations between airborne microorganisms and meteorological conditions, and the influence of local versus long-range transport of microbes, air samples were continuously taken from a platform at the 458-m level of the Tokyo Skytree (a 634-m-high broadcasting tower in Tokyo) from August 2016 to February 2017. Predicted source regions of airborne particles, from back trajectory analyses, changed abruptly from the Pacific Ocean to the Eurasian Continent in the beginning of October. However, microbial community composition and alpha and beta diversities were not affected by this meteorological regime shift, suggesting that long-range transport from ocean or continent was not the principal determinant controlling the local airborne microbiome. By contrast, local meteorology, especially relative humidity and wind speed, had significant relationships with both alpha diversities and beta diversity. Among four potential local source categories (soil, bay seawater, river, and pond), bay seawater and soil were constant and predominant sources. Statistical analyses suggest humidity is the most influential meteorological factor, most likely because it is correlated with soil moisture and hence negatively correlated with the dispersal of particles from the land surface.

## Introduction

Microorganisms are abundant in the atmosphere, with numbers ranging from 10^4^-10^5^ cells m^−3^ in ambient air on mountain peaks [1] to 10^6^-10^7^ m^−3^ in desert dust storms [2]. Air is now recognized as a highly diverse microbiome [1, 3-14]. Airborne bacteria and fungi also have the potential to cause human diseases [15]. For example, airborne fungi have been identified as the cause of respiratory problems such as asthma after thunderstorms [16], and also the lymph node syndrome, Kawasaki disease [17]. Endotoxins from airborne bacteria are also associated with health issues [18]. Airborne microorganisms are important not only for human health, but also for climate and ecology. For example, the plant pathogen *Pseudomonas syringae*, and related phylloplane bacteria, have strong ice nucleation ability at very warm temperatures (>-5 °C, 19-22) and airborne ice nucleating bacteria may promote the formation of ice in clouds, which may modify their radiative forcing and promote precipitation. Until recently, most airborne microorganism studies depended on conventional culture-based methods [e.g. 23, 24]. However, culture-independent methods using Next Generation Sequencing (NGS) are now prevalent [1, 3-14]. This technology transforms our ability to describe spatial and temporal variability of all airborne microorganisms, and enables us to compare their temporal variation under different meteorological conditions. However, the relationship between aerial microbial community composition and meteorology is still not well characterized [25]. For example, Bowers and co-workers [5] showed different land-use types, rather than local weather, control the airborne bacterial composition in northern Colorado, US during the early summer season. On the other hand, Gandolfi and co-workers [9] showed wind speed and relative humidity affected airborne microbial community structure in two urban sites, in northern Italy. Šantl-Temkiv and co-workers [11] showed that the alpha diversity of bacterial communities is positively correlated with air temperature and negatively correlated with relative humidity in western Greenland during mid-summer.

Studies are often focused on urban areas, because these contain potential sources of human pathogens, such as water treatment facilities and densely populated areas. While spatial and temporal variations of bioaerosols in urban environments have been frequently studied in many cities [8-10, 13, 26, 27], few studies [6] have used consecutive short-period samples to explore correlations between variation of the microbiota and meteorology. Additionally, samples from these studies were taken near ground level (generally several meters above) or on roof tops of lower buildings, with the result that specific local sources very near to sampling sites are likely to predominate over microorganisms originating regionally or from afar, and so obscure the effect of remote contributions.

Tokyo (population: 13.61 million) is located south-east of the Japanese main island. The climate of Tokyo is warm and humid in summer and cool and dry in winter. Despite the city’s size, bioaerosol studies of the capital are limited to a recent study of bacterial community composition in rainfall [28] and a study of bacteria in railway stations [29]. In order to investigate 1) airborne bacterial communities over Tokyo, 2) fine-scale correlations between airborne microorganisms and meteorological conditions, and 3) the influence of local versus long-range transport of microbes, we obtained a series of consecutive 48-72 h samples at a height of 458 m from the tallest communication/observation tower in Japan, the second tallest structure in the world (Tokyo Skytree: 634 m), from summer in 2016 to winter in 2017.

## Method

### Sampling of air and reference samples

Airborne microorganisms were collected at a 458-m level measurement site [30] on the western side of the Tokyo Skytree (SI Table 1) from August 2016 to February 2017. Eight sets of 2 m length conductive silicon tubing were fixed to the outside wall of the Tokyo Skytree with their inlets under a wind and rain shield, and sterilized inline NILU filter holders (Norwegian Institute for Air Research) attached to the inner ends. These were fitted with pre-combusted (500 °C for 2 h) 47 mm diameter quartz filters (Tissuquartz™ Filters, 2500 QAT-UP, Pall). The vacuum sides of the NILU filter holders were connected to low volume samplers (LV-40BW, Sibata Scientific Technology) programmed to sample from one unit at a time. Flow rate was 15-30 L min^−1^ for 48-72 h for each filter, and then the sampler switched to a new unit. Sampling was continuous. Sampling details are described in SI Table 2. In order to avoid degradation of filter samples during and after sampling, all filter holders were placed in a 4 ℃ refrigerator (JR-N40G, Haier) until retrieval (usually at two – three week intervals). Reference environmental (potential source) samples for source estimation were taken around Tokyo Skytree on Jan 3, 2017 and from around the building of the National Institute of Polar Research (NIPR) on June 6, 2016 (SI Table 1, SI Fig. 1). Soil samples were placed in a 5 mL sterile plastic tube containing 2 mL of RNAlater (Thermo Fisher Scientific, MA, USA) using a pre-cleaned stainless-steel spoon. Seawater, river water and pond water were sampled directly into a 50 mL sterile plastic bottle. All soil and water samples were kept at −20 °C prior to DNA extraction.

**Fig. 1:**
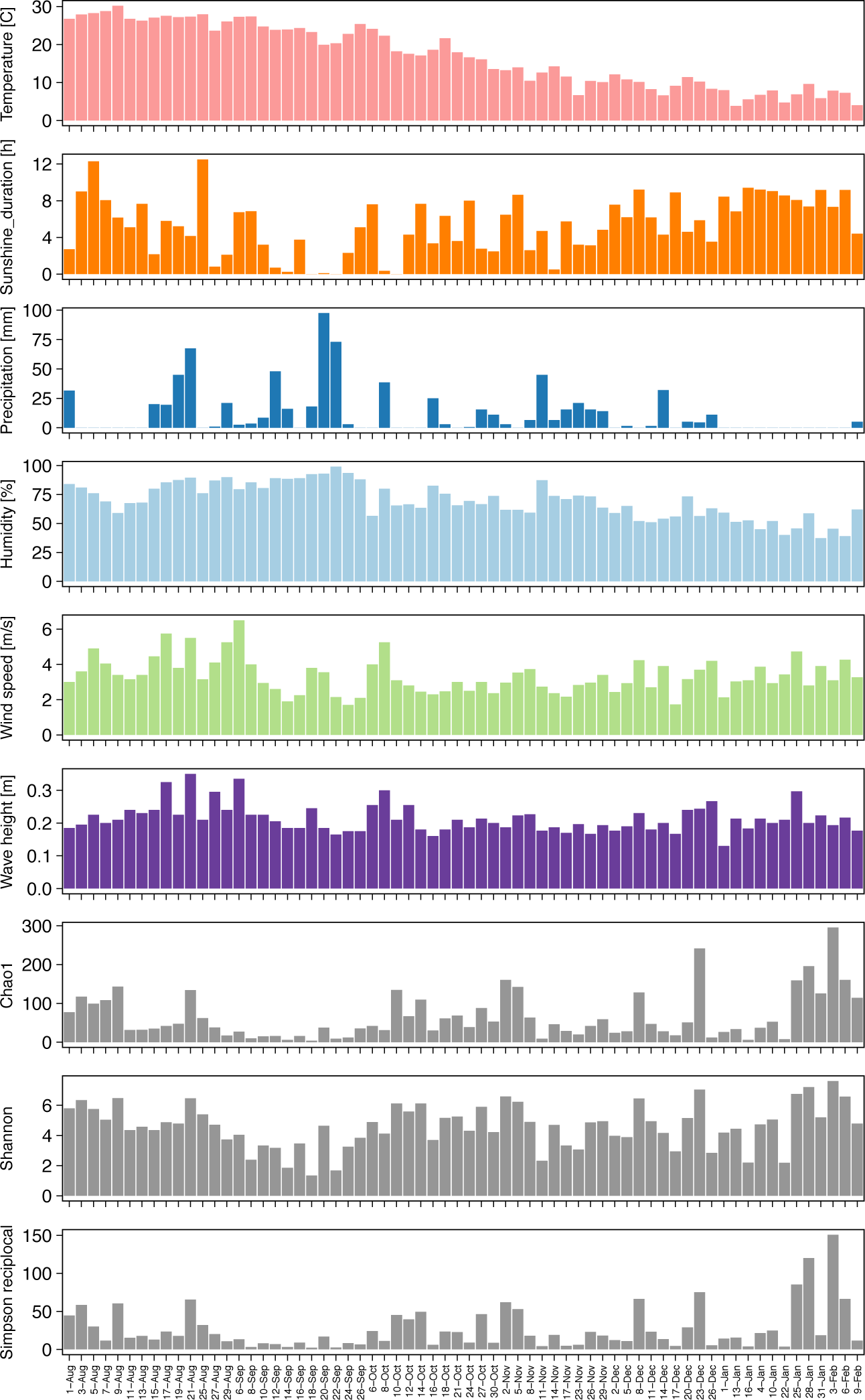
Seasonal changes of meteorological factors (temperature: pink, Sunshine duration: orange, precipitation: blue, humidity: light blue and wind speed: light green) from nearest automatic weather stations, wave height (purple) and alpha diversities (gray) during sampling periods.

### DNA extraction, PCR and DNA sequencing

In order to avoid contamination, all processes prior to Polymerase Chain Reaction (PCR) amplification were done in a laminar flow clean bench (PCV-1305BNG3-AG, Hitachi). The clean bench was sanitized with a UV lamp overnight, and pipettes were sterilized in a DNA cross linker (CL-1000, UVP) box inside the clean bench. Genomic DNA in bioaerosols captured on quartz filters was extracted using the FastDNA™ SPIN Kit for Soil (MP Biomedicals, Santa Ana, CA). The quartz filter was initially pulverized during the bead beating step, but in order to maximize the yield of DNA (DNA adsorbs to quartz fibers), all fragments of the filter were carried over until the final elution step. Partial 16S rRNA gene sequences, spanning the V3 and V4 regions, were amplified using the primers Bakt_341F (5’ CCTACGGGNGGCWGCAG 3’) and Bakt_805R (5’ GACTACHVGGGTATCTAATCC 3’), with Illumina overhang adaptor sequences attached to their 5′ ends, by KAPA HiFi HotStart ReadyMix (KAPA Biosystems, MA, USA). PCR reaction conditions comprised 35 cycles of denaturation at 95 °C for 30 s, annealing at 55 °C for 30 s, and elongation at 72 °C for 30 s and an additional final elongation at 72 °C for 5 min using a GeneAmp PCR System 9700 (Applied Biosystems, CA, USA). Subsequent clean-up and indexing of PCR amplicons were performed following Illumina methods for 16S metagenomic sequencing library preparation (https://support.illumina.com/content/dam/illumina-support/documents/documentation/chemistry_documentation/16s/16s-metagenomic-library-prep-guide-15044223-b.pdf). All samples were pooled into one flow cell of a MiSeq sequencer (Illumina, San Diego, CA) and sequenced at NIPR using a MiSeq V3 reagent kit.

### Exact sequence variant analysis

In order to understand sequence differences in greater detail than the conventional 97% operational taxonomic unit (OTU) approach, we used a newly developed DADA2 (1.4) method which, by incorporating an error model, is able to infer sequence reads with single nucleotide resolution [31]. All air samples, negative PCR amplicons, reference samples and potential contamination from previously analysed glacier samples in NIPR (DRA004099– DRA004103) were analysed together with filtering parameters: truncLen=c(250,210), maxN=0, maxEE=c(1,3), truncQ=2. Taxonomy was assigned by the Naive Bayes Classifier method in The Ribosomal Database Project (RDP) Classifier [32] implemented in the DADA2 program against the Silva 128 database [33]. All potential chimera sequences were removed by DADA2 and all chloroplast and mitochondria were removed manually. Alpha diversities, rarefaction curve of alpha diversities, unweighted UniFrac distances analysis [34] and unweighted Pair Group Method with Arithmetic Means (UPGMA) were analyzed by QIIME [35]. Based on rarefaction curve results and number of filtered sequences, sequences are rarefied to 970 for alpha diversity and 1000 for beta diversity estimation. A taxonomy heatmap was created by the R Package “superheat” [36]. Sequence data (DRA007168-007176) are available from DDBJ Sequence Read Archive (DRA): https://www.ddbj.nig.ac.jp/dra/index-e.html.

### Source estimation

“SourceTracker” is a Bayesian approach program to estimate the proportion of exogenous sequences in a given community that come from possible source environments [37]. We used the latest version, SourceTracker2 (https://github.com/biota/sourcetracker2), for source estimation of bioaerosols sampled. Twenty three reference samples collected around Tokyo Skytree (8 sites: SI Fig. 1) were analyzed as possible sources of airborne bacteria. In order to focus on local bacteria around the Tokyo area, 6 Amplicon Sequence Variants (ASVs), which have significant differences (p<0.001, Mann-Whitney) between non-continental and continental periods, were removed from analysis.

### Statistical data analysis

Correlation between alpha diversities, source contributions, and meteorological data were analyzed by Spearman’s correlation using the R package “ggcorrplot”. Mann-Whitney U tests between non-continental and continental periods were performed with XLSTAT software (https://www.xlstat.com). Analysis of similarities (ANOSIM) and permutational multivariate analysis of variance (PERMANOVA) for unweighted UniFrac dissimilarity matrices were performed using PRIMER 7 (Plymouth, UK).

### Meteorological data

All meteorological data taken by the Automated Meteorological Data Acquisition System (AMeDAS), which is managed by the Japan Meteorological Agency (JMA), are available from http://www.data.jma.go.jp/obd/stats/etrn/index.php. Air temperature, wind speed, sunshine duration (total duration of solar radiation above 0.12kW m^−2^) and humidity are taken at the “Edogawa rinkai” station and precipitation measures were taken at the “Tokyo” station (SI Table 1). Wave height data in Tokyo Bay, which is managed by the Bureau of Port and Harbor, Tokyo Metropolitan Government, are available from http://www.kouwan.metro.tokyo.jp/yakuwari/choui/kako1-index.html. All daily data were transformed to 2-3 day averages to correspond to sampling durations (SI table 2, Fig. 1). The trajectories of air masses arriving at the sampling location were calculated by HYSPLIT (https://ready.arl.noaa.gov/HYSPLIT.php [38]) using GDAS 0.5 model (lat = 35.7101, lon = 139.8107, height = 458, duration = 72) and were started from 12:00 (UTC) of each day (Fig. 2). All samples were categorized as either continental, meaning of Eurasian continent origin, or non-continental, based on their latitude/longitude location 72 h prior to arriving at Skytree.

**Fig. 2:**
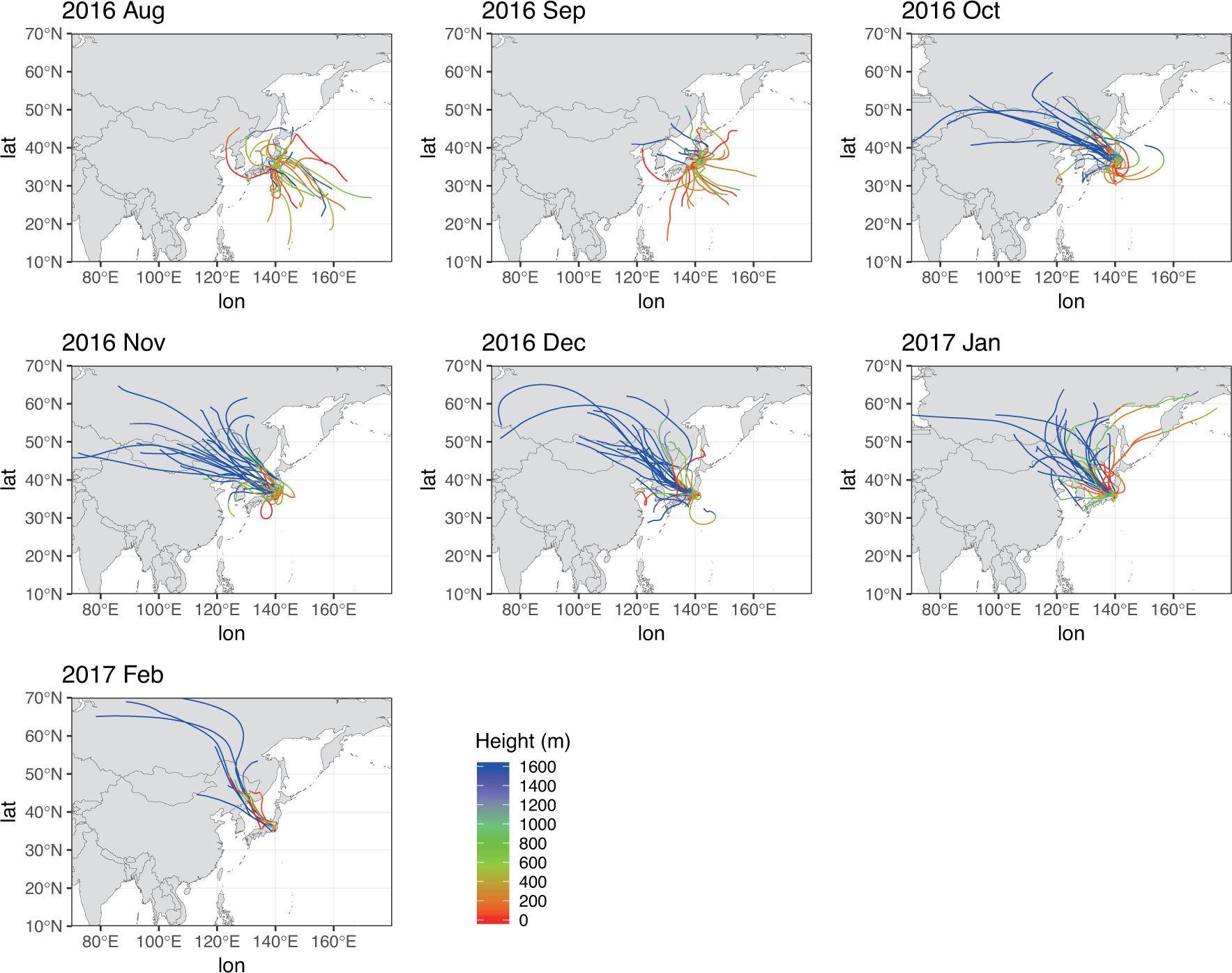
72h HYSPLIT back trajectories during sampling period. Height above 1600 m is shown in blue.

## Results

### Taxonomy

An average of 5862 sequences per sample (maximum: 19913, minimum: 1068) were used for analyses (SI Table 2). Figure 3 shows temporal variation of the major phyla. *Proteobacteria* (mean 51.4% of relative abundances) was by far the most common phylum followed by *Firmicutes* (13.6%), *Cyanobacteria* (7.9%), *Actinobacteria* (7.7%), *Bacteroidetes* (5.2%), and *Acidobacteria* (3.0%). Of all the phyla, only *Parcubacteria* (2.6%) exhibited seasonal changes, being higher during August and September. Finer resolution of taxonomy is shown in SI Figure 2, which gives the temporal variation of the 100 most abundant major genera. *Paraburkholderia* (mean 6.3%) had the highest mean relative abundance throughout the entire sampling period, followed by *Sphingomonas* (3.5%), *Chroococcidiopsis* (3.3%), *Bacillus* (3.2%), *Pseudomonas* (3.1%), *Stenotrophomonas* (2.8%), *Methylobacterium* (2.7%), *Craurococcus* (2.2%), an unidentified genus in order *Rhizobiales* (1.9%), *Mesorhizobium* (1.8%), *Clostridium* (1.6%), and an unidentified genus in the class *Adlerbacteria* (1.4%). Relative abundances of some genera were significantly (p<0.05, Mann-Whitney) different in early October (SI Table3), when the HYSPLIT model indicated that the source region was shifted from non-continental to continental between October 2^nd^ and 4^th^.

**Fig.3:**
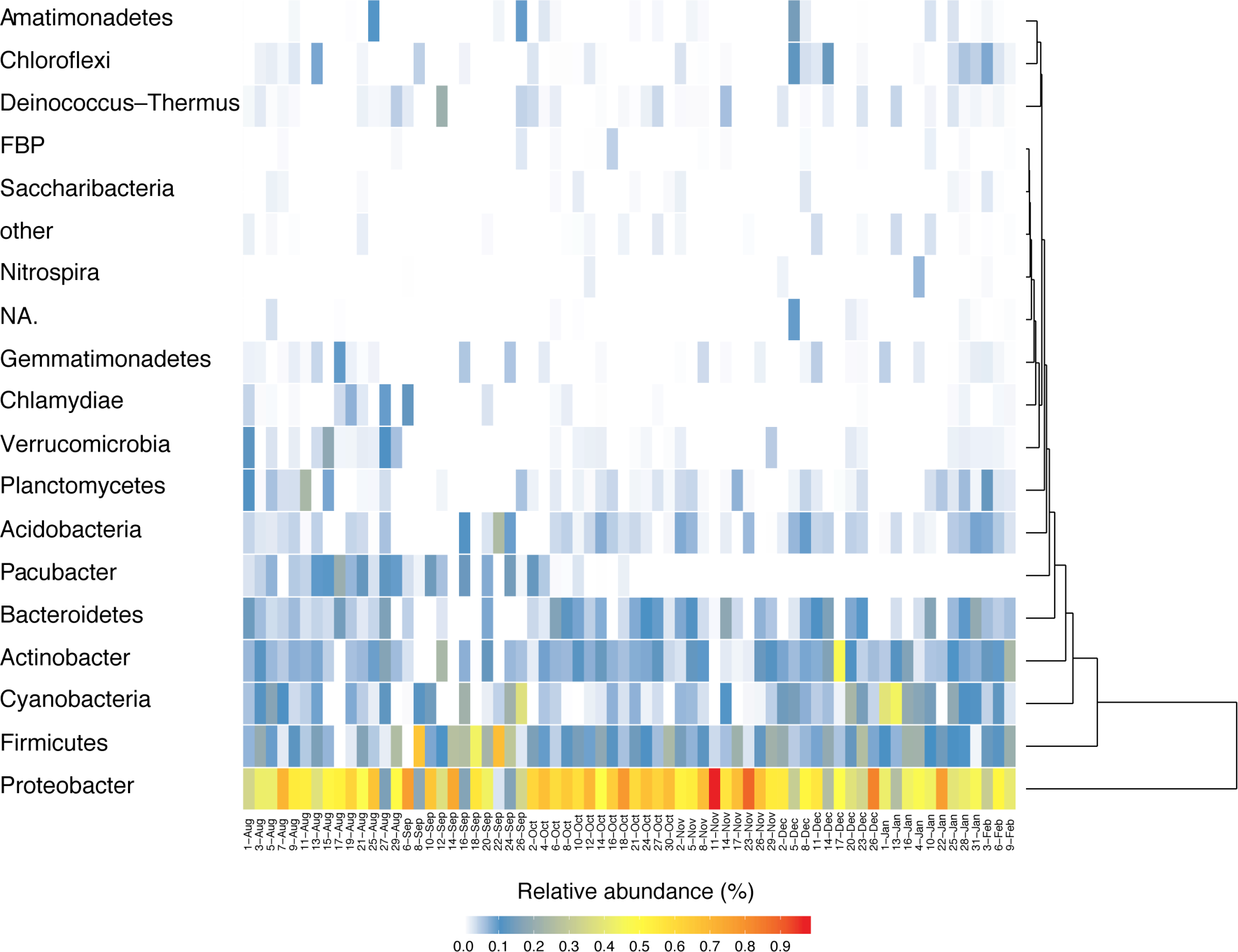
Seasonal change of bacteria at the phylum level.

Six ASVs (mostly members of the *Parcubacteria*) out of a total of 3257 ASVs had significant differences in their abundances (p<0.001, Mann-Whitney) between non-continental and continental periods (SI Table4, SI Fig. 3). MetametaDB analysis shows that the estimated sources of five of these ASVs (all the *Parcubacteria* and an *Opitutus* sp. from the *Verrucomicrobia*), that were abundant in the non-continental period, are from freshwater environments. The sixth ASV (*Paraburkholderia* sp.) was abundant in the continental period, and is mostly from soils (SI Fig. 4). The closest matches using BLAST searches with these 6 ASVs confirmed the same potential sources (SI Table 4). Correlations between all ASVs (excluding the 6 ASVs mentioned above) and the 6 local meteorological and ocean factors were significant (correlation coefficient >±0.4, p<0.05) only for a *Sphingomonas* sp. (ASV714) with temperature (0.46, P = 0.0001); ASV714 matched with 100% identity to sequences from rainfall (KX508562), urban aerosol (DQ129617) and plant surfaces (AB582140, HM450020).

**Fig. 4:**
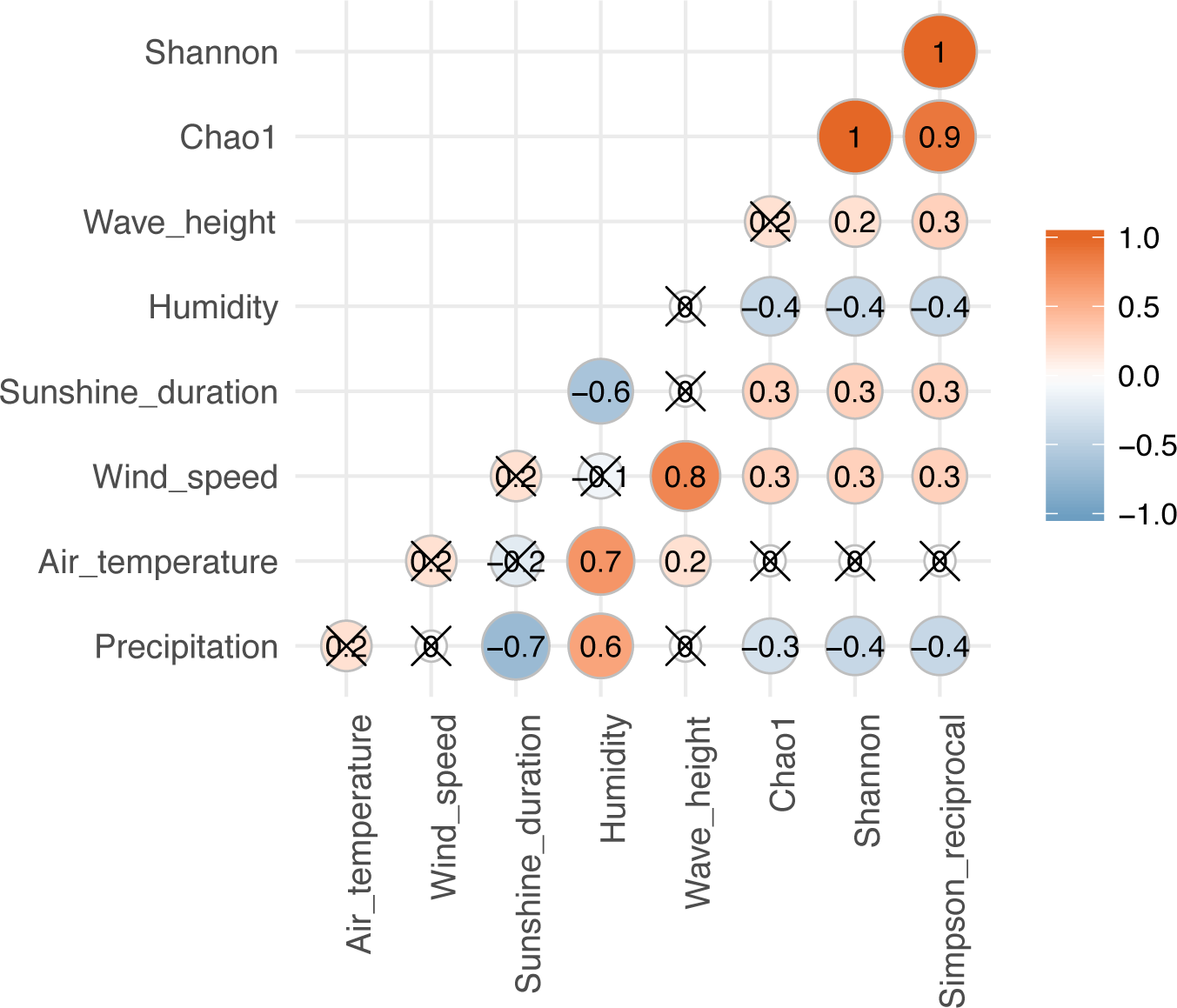
Spearman’s correlation between alpha diversities and meteorological and ocean factors. A red/orange circle shows positive correlation and a blue circle shows negative correlation. A cross “X” on the circle indicates no significance (p < 0.05).

### Alpha diversity

Three alpha diversity methods were measured: Chao 1, the Shannon index (H’), and the reciprocal of Simpson’s index (D’). Chao 1 estimates total ASV richness, the Shannon index is a general diversity measure that is positively correlated with both diversity and evenness, being sensitive to differences in abundance of rare ASVs, and Simpson’s reciprocal is a measure of evenness, which has a lower bound of 1 for a community composed of only one ASV and, for example, a value of 10 for a community containing 10 equally abundant ASVs [39]. Adequate coverage is required for reliable estimation by the indices [39], and indeed the depth of sequencing was enough for all three to stabilize (by 1000 sequences; SI Fig. 5). Obvious transitory peaks in alpha diversities occurred in August, October, and the end of January and at the beginning of February (SI Table 1 and Fig.1). There was also and extended period of consistently low alpha diversity during September. Correlation coefficients with the 6 local meteorological and ocean factors are shown in Fig. 4. Among meteorological and ocean data, humidity and precipitation had the most significant negative correlations with all diversity measures, while wind speed and sunshine duration had significant positive correlations with the alpha diversities.

**Fig. 5:**
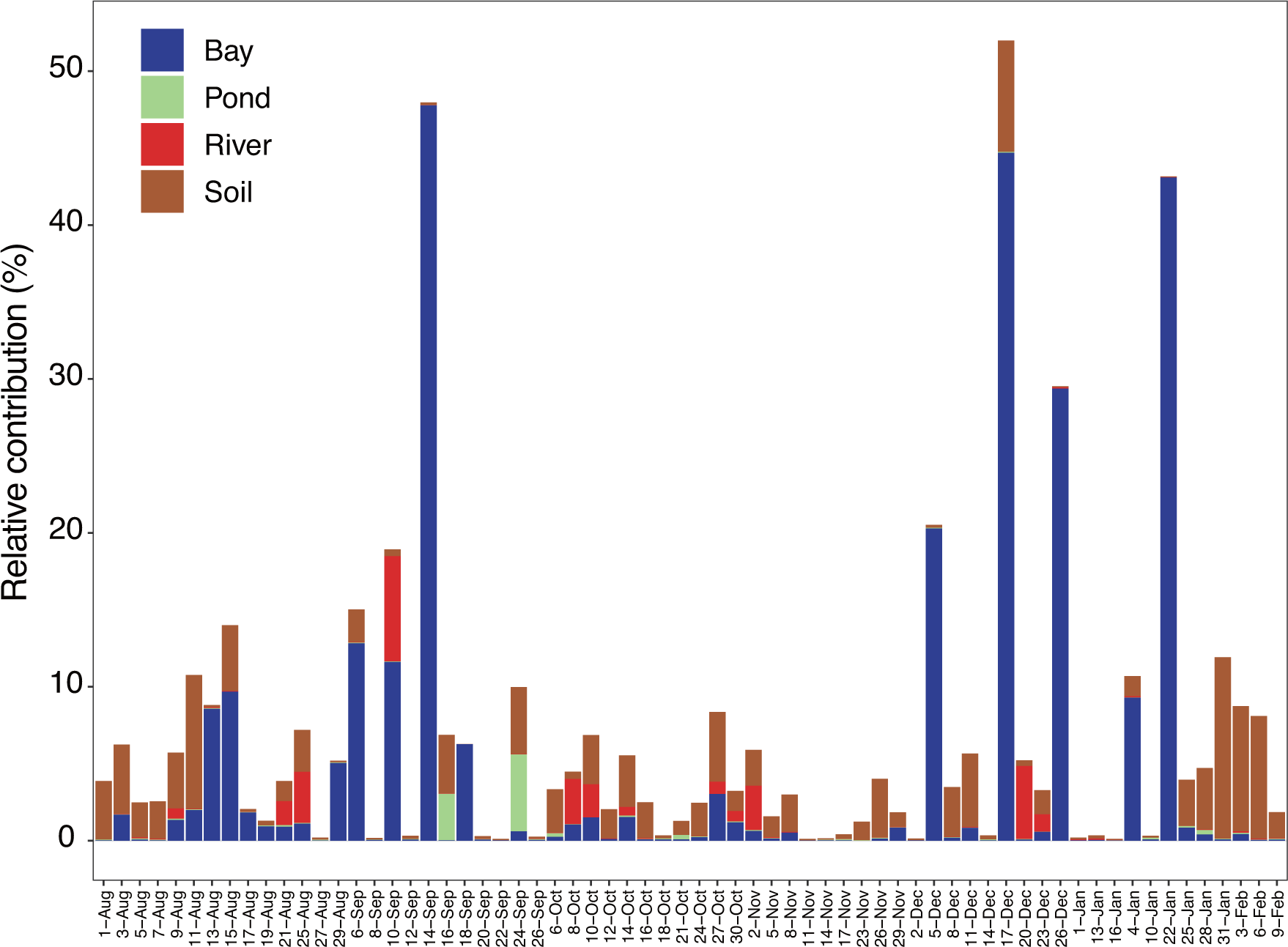
Seasonal change of estimated contribution from potential source types (bay, soil, pond and river) by source tracking analysis.

### Beta diversity

A unweighted UniFrac dendrogram was clustered into two groups with a Bray-Curtis dissimilarity of 0.86, with a lower alpha diversity cluster and higher alpha diversity cluster (SI Fig. 6). ANOSIM, analyzed by differences in HYSPLIT source, month and temperature, shows higher ANOSIM R (0.337 for HYSPLIT, 0.396 for month, 0.274 for temperature) in the higher alpha diversity cluster (SI Table 5). A PERMANOVA test on the effect of meteorological data showed humidity and wind speed were strongly related to unweighted UniFrac distance (PERMANOVA, p <0.005, SI Table 6).

**Fig. 6:**
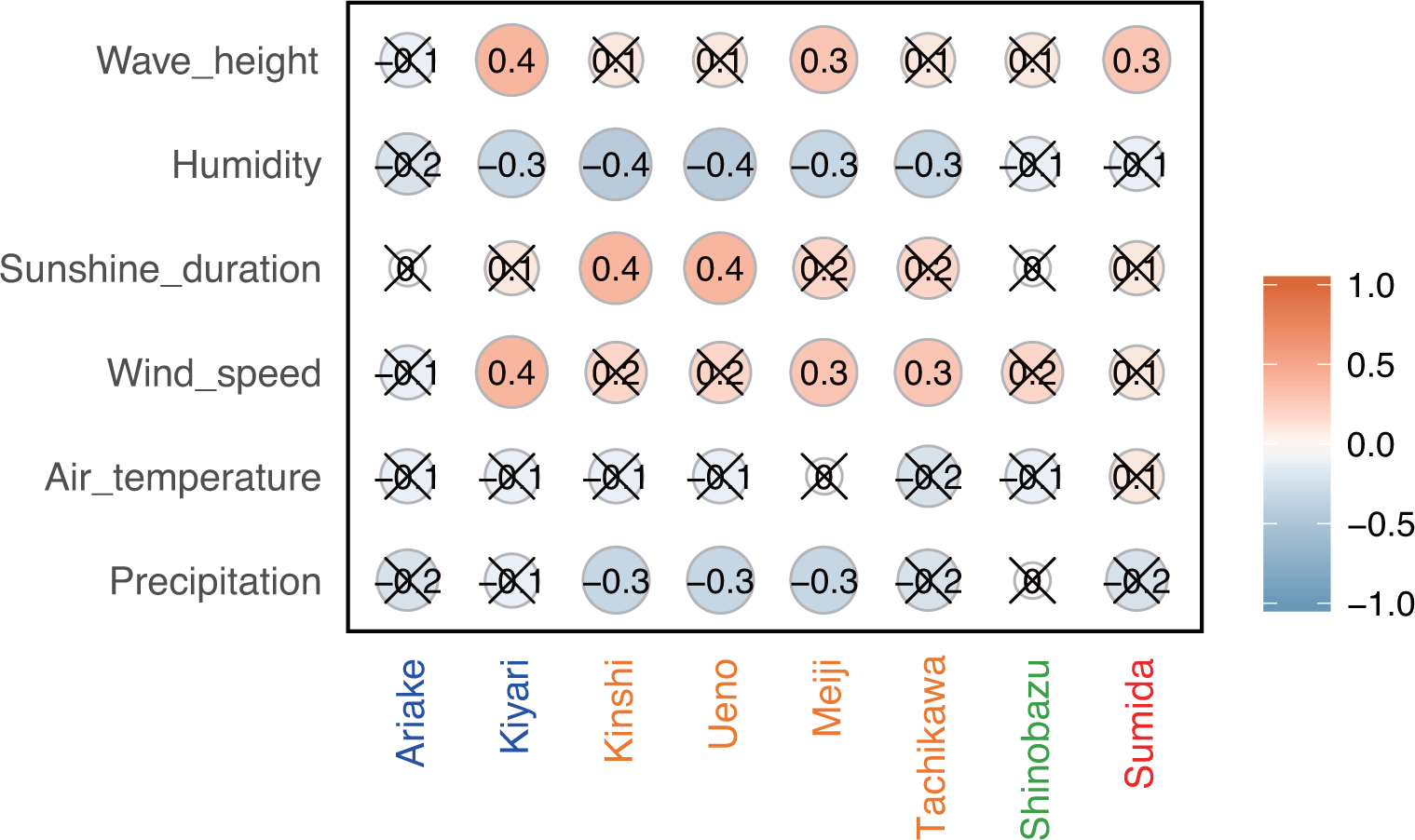
Spearman’s correlation between estimated source contribution and meteorological factors and wave height in Tokyo Bay. A red/orange circle shows positive correlation and a blue circle shows negative correlation. Cross “X” on the circle indicates no significance (p < 0.05).

### Source estimation

Sourcetracker2 analysis shows the estimated contributions to Tokyo Skytree air samples from each potential source (Fig. 5 for each category, SI Fig.7). Total percentage contributions from the tested sources had a maximum of 52%, a minimum SI of 0.1%, and a mean of 6.8%. Bay influence was relatively higher in the early sampling period and occasionally of influence at later periods. Soil is the second highest contributor, contributing throughout the sampling period. Correlations between source contributions and environmental factors are shown in Fig. 6. Among the six environmental factors, humidity had a significant negative correlation for all soil sites, while wind speed had significant positive correlations for three soil sites (Kiyari, Meiji and Tachikawa).

## Discussion

### Long-range effects of air mass shifts upon bacterial composition

Bacteria, which have typical diameters of ∼1µm, are readily emitted from surfaces under certain conditions, and have aerial residence times from days to months [40, 41], during which time they may be transported over great distances before their removal by both wet and dry deposition [42-44]. On non-dusty days, bacterial concentrations near ground level (10 m a.g.l.) are much higher than in air at higher levels [44], indicating that near-ground levels are strongly influenced by local sources. The average daytime atmospheric boundary layer height generally exceeds 500 m, even in wintertime, but is generally just below 500 m at night time over all seasons [45]. Therefore, the Tokyo Skytree sampling site will sample boundary layer air primarily during daytime, and air from the residual layer and potentially the lower free troposphere during nighttime. Since samples were obtained continuously over several days, the deposited particulate matter in any one sample will be derived from this mix of air masses. ANOSIM analyses on unweighted UniFrac distances clearly clustered into non-continental and continental sources (ANOSIM Global R =0.337, P < 0.001), as predicted using HYSPLIT back trajectories. This finding is similar to a previous study showing strong seasonal patterns (ANOSIM Global R =0.47, P < 0.001 from [4]). At the genus level of taxonomy, the relative abundances of *Paraburkholderia* and an unidentified genus in the phylum *Parcubacteria* are significantly changed between non-continental and continental sources, respectively (SI Fig. 2). Therefore, these genera are presumed to have originated from non-local sources, indicating that synoptic air mass movement influenced bacterial distributions.

To obtain more detail on these potentially long range transported bacteria, we investigated their ASV level taxonomy. Using this, only 6 ASVs out of 3257 ASVs were identified as potential long range transported bacteria on the basis of trajectory analysis. These were relatively abundant, especially ASV137, which belongs to the genus *Paraburkholderia*, with notably high relative abundance in the later sampling period (SI Fig. 3). Nevertheless, the majority of ASVs were not associated with long range air mass movement. This implies that most are of local origin, and therefore were influenced by local meteorological conditions. We focus discussion in this section on the six ASVs of potential long-range sources.

Five of the six ASVs, belonging to the genus *Parcubacteria* (ASV429, 609, 1233, 1036) and *Opitutu* (ASV 1556), were expected to be pelagic because they were abundant only during air mass flows from over the Pacific Ocean. However, these are likely to be freshwater bacteria according to their closest matches using BLAST searches and metametaDB analysis. Their non-oceanic origin is also supported by the absence of ocean bacteria, such as bacteria belonging to the SAR group or typical marine groups, such as *Oceanospirillales*, *Spongiibacteraceae* and *Marinicella*. Thus, despite the HYSPLIT results, this composition indicates minimal inputs from the Pacific Ocean. This can be explained by a combination of low emissions from the ocean surface and higher inputs from regional freshwater. In previous studies, similarly lower contributions of ocean bacteria were reported. For example, the number of cultivable bacteria found in near-surface air during non-dust events over the ocean is generally very low [42, 46], and the number of marine bacteria in the near-surface air of a coastal city in Greece was found to be rare [10].

One of the six ASVs, belonging to the genus *Paraburkholderia* (ASV137) and abundant in the continental period, was indicated to be a soil bacterium based on closest relative matches in BLAST searches and metametaDB analysis. Exactly same sequences were found from soil in many countries including China (e.g. KU323602, KX351056, KY427125). Since these were transported by westerly winds, which prevailed in the later sampling period when air masses mostly came from the Eurasian Continent, this ASV is possibly from Chinese soil. Although some bacteria are known to be transported from arid regions in China during Asian dust events in March and April in Japan [47], the genus *Paraburkholderia* (and its higher taxonomy order *Burkholderiales*) was not found in ground observations during Asian dust days [48], in the upper Asian dust layer [44] or from the Gobi Desert, one of their potential source areas [2]. Therefore, ASV137 is not likely to be of Asian dust origin. Furthermore, *Bacillus subtilis*, which is common in Asian dust studies [44, 49], was found only once over the entire sampling period. Some *Bacillus* ASVs are relatively close to uncultured bacteria from the Taklimakan Desert (AB696509 and AB696498), however, these were not perfectly matched by BLAST (ASV399: 99.1%, ASV6088: 98.6%, ASV1582: 98.1%). Therefore, we expect that the soil type, ASV137 inhabited, is not desert type soil.

### Local meteorological effects upon sources of bacteria

Most ASVs were associated with airmasses arriving from local regions prior to sampling, therefore, it is most appropriate to investigate relations of these bacteria with local meteorological conditions. Local meteorology, especially relative humidity and wind speed, had significant relationships with three alpha diversities (by Spearman correlation) and beta diversity (by PERMANOVA). While previous studies failed to analyze the cross-correlated meteorological variables [41], multivariate statistics enable these relationships to be revealed [6]. We found that both simple correlations and multivariate analyses showed the same relationships, with humidity and wind speed being effective meteorological predictors of bacterial composition and diversity.

The effect of relative humidity change upon airborne bacterial community composition was reported in some prior studies. For example, canonical correspondence analyses of bacterial community structure in urban bioaerosols in Italy showed both relative humidity and wind speed were variables affecting airborne bacterial community structure [9], and a study in western Oregon, U.S.A showed that airborne bacterial concentrations were positively correlated with temperature but negatively correlated with relative humidity [50]. In the present study, relative humidity increased during precipitation events and remained high, before decreasing with a sufficient amount of radiation (Fig 1). Relative humidity is especially high in late August – September, when it was strongly affected by a passing typhoon and an autumn rain event; alpha diversities were, appropriately, low in the same period. High humidity tends to keep the surface of the ground wet, and the bonding force by surface tension will keep the particles attached to the surface. This bonding force will be reduced by drying [11, 51]. Accordingly, among the three different source types, contributions from all the soils (Kiyari, Ueno, Meiji and Tachikawa) had significant negative correlations with relative humidity, with contributions higher in the middle of winter, the driest season during the sampling period (Fig. 1, Fig. 6).

Precipitation, which obviously strongly affects the humidity and surface bonding forces, was not found to have a consistent or significant relationship with airborne bacterial diversity and composition. While alpha diversities tended to decrease when precipitation occurred (SI Fig 8: blue broken lines), exceptions occurred during heavy precipitation events resulting from a typhoon that set historical precipitation records and during some stationary fronts (SI Fig 8: red broken lines). The common expectation, and what occurred during the majority of lighter precipitation events, is that airborne bacterial communities in Tokyo would be reduced by precipitation due to scavenging (wet deposition) [49, 52] and wetting of the soil surface. However, alpha diversities slightly increased during the historical heavy rains on Aug. 21-22, Sep. 12-13, and Sep. 22-23 (SI Fig. 8: red broken line). For example, rain rates of 107.5mm/h were reported in Ome (30km west from site) on Aug. 22 during typhoon passage, and 63.5mm/h in Yokoshibahikari (80km east) under a stationary front condition. Diversity increase during such heavy rain events may be explained by bioaerosol generation caused by the impact of rain drops on plant, soil and built surfaces [50, 53, 54], and in this case potential sources would be near to the sampling site (e.g. even the outside walkway and wall of the tower).

Wind speed has been shown to be positively correlated with airborne microbial concentrations in some previous studies [55-57]. As well, active water surfaces enhance aerosol generation into the atmosphere [7, 44, 58-60]. Wind speed is highest in late August, and was significantly correlated with wave height in Tokyo Bay (Fig. 1). High wind speeds over the bay’s surface resulted in rougher waters, which would be expected to correlate with more aerosol generation during wave-breaking (via the bubble bursting process) [41]. Among the three potential sources, the bay sample Kiyari had the most significant positive correlation of source contribution with wind speed and wave height. Wind speed was also weakly correlated with some of the soil sites (Meiji and Tachikawa), suggesting that dispersal of bacteria from the soil surface is enhanced during stronger winds.

Many previous studies showed temperature, which follows seasonal cycles in temperate regions, is also a factor controlling airborne bacterial emissions and community composition [6, 10, 50, 51]. In contrast, temperature was not found to be a significant factor in this study. The effects of local meteorological conditions upon airborne microorganisms is still controversial, because recent studies have suggested that airborne bacterial communities could be more affected by variation in local sources than by changes in local meteorological conditions [5, 6]. Our findings oppose these studies, as also observed in other work on the effects of raised relative humidity and rainfall upon primary bioaerosols and ice nucleating particles [61-65]. However, the relationship between bacteria and meteorological factors undoubtedly varies between different geographical locations and ecotypes.

In this study, air mass shifts detected by HYSPLIT analysis indicated limited long-range effects. Estimated contributions from local sources, from seawater in Tokyo Bay and soil, were consistently found and predominated over other known sources over the sampling period (Fig. 5). Since the samples for source estimation were limited in type, number of sites, and season, our reference samples only explained, on average, < 6.9% of the total airborne bacterial communities. This implies that we should include more diverse potential source types in future studies, such as plant surfaces [66] and human impacts in highly populated areas [6]. These improvements will likely enable more accurate and comprehensive source estimations. Finally, although we took samples from the highest structure in Japan, it was still primarily within the urban atmospheric boundary layer. If comparisons were made with samples from above the boundary layer, we expect that the ratio of long-range-transported bacteria would be much higher.

In order to obtain enough DNA for analysis, we obtained samples that were integrated over 2-3 days. However, shorter interval sampling may be helpful for understanding the effect of radical meteorological changes upon microbial communities, such as heavy rain enhancement of microbial emissions versus precipitation scavenging in other rain events. Recently, much higher flow rate and lower cost sampling devices have been proposed [67] for use in the field [11]. Such capabilities may offer the ability to understand not only hourly changes of specific microbes, but also offer sufficient sample sizes to permit metagenomic analyses of potential microbiome function.

## Conclusions

Consecutive 48-72 h samples from the tallest tower in Japan, combined with NGS analyses, were used to reveal how airborne microbial communities changed from summer to winter. In this study, air mass shifts detected by HYSPLIT analysis indicated limited long-range effects on microbial populations. Our results showed only a limited number of ASVs that could potentially be associated with long distance transport of bacteria. However, most of the ASVs were not affected by abrupt air mass changes, indicating they were likely to have been from local sources. Local inputs, from soil and the seawater in Tokyo Bay, were consistently found and the predominant sources over the study period. Humidity and wind speed were key factors affecting bacterial alpha and beta diversity, and hence, these appear to be the controlling factors on emissions of bacteria from bay sea water and soil around the Tokyo Skytree. NGS allowed us to study the airborne microbiome with unprecedented resolution, and the accumulation of such knowledge from many environments across the world [1, 3-14] could provide a more comprehensive understanding of factors determining local, regional and global airborne microbiomes.

## Acknowledgements

The authors thank to Mr. Kenichi Watanabe, Ms. Ayumi Akiyoshi and Ms. Mizuho Mori in for assistance in laboratory experiments. This study was partially supported by crowdfunding operated by academist Co., Ltd. The authors specially thank all supporters of Jun Uetake’s project (https://academist-cf.com/projects/36). Jun Uetake was supported for further analyses of data under National Science Foundation grant AGS1660486. The authors gratefully acknowledge the NOAA Air Resources Laboratory (ARL) for the provision of the HYSPLIT transport and dispersion model and the READY website (https://www.ready.noaa.gov/index.php) used in this publication.

