## Supplemental Figures for "Seasonal changes of airborne bacterial communities over Tokyo and influence of local meteorology"

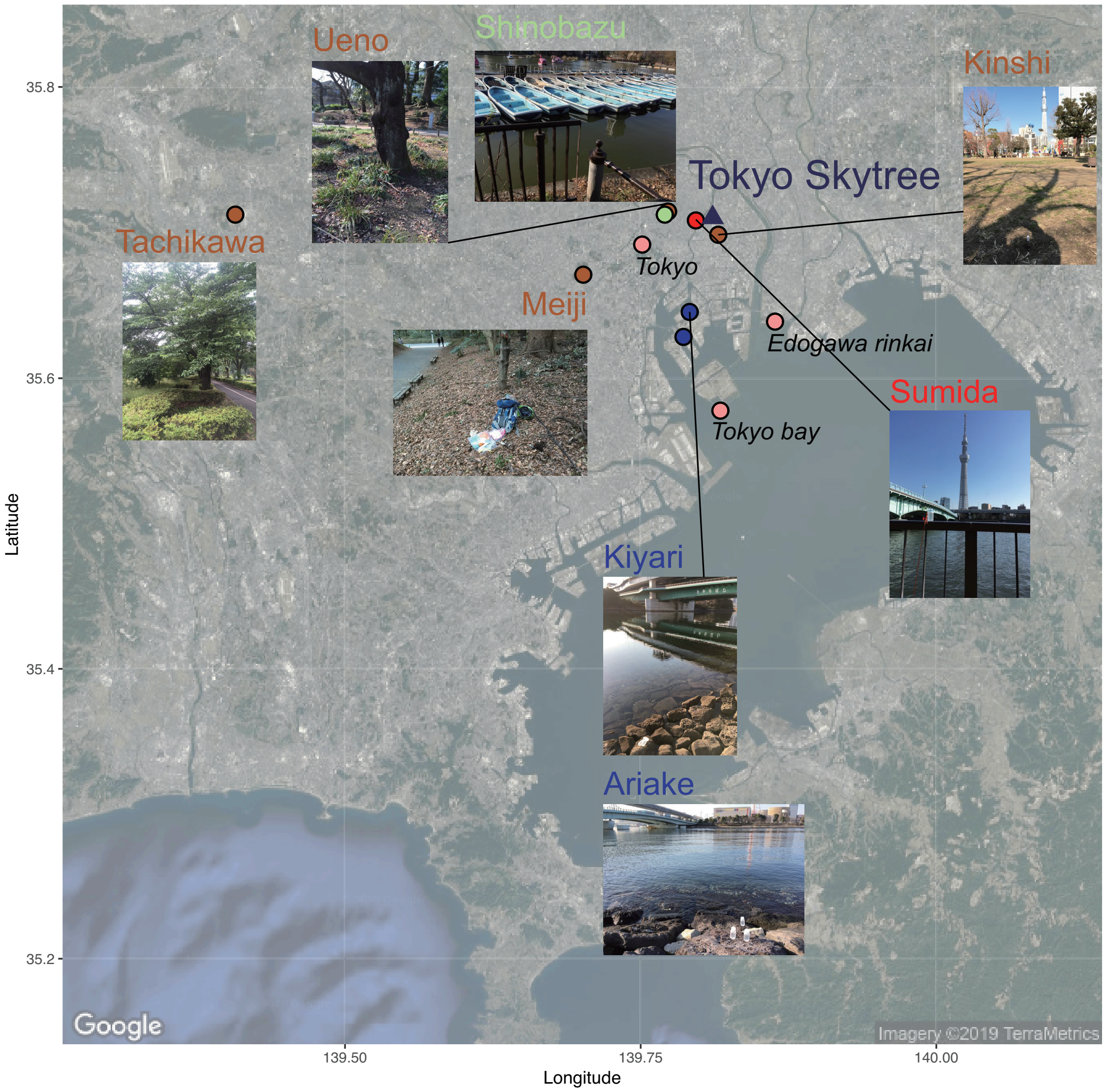


SI Fig. 1: Location of Tokyo Skytree (blue triangle), sampling sites for reference (blue circle: bay, brow circle: soil, red circle: river and green circle: pond) and meteorological and oceanic observation sites (pink circle). More detailed information are available in SI Table.1.

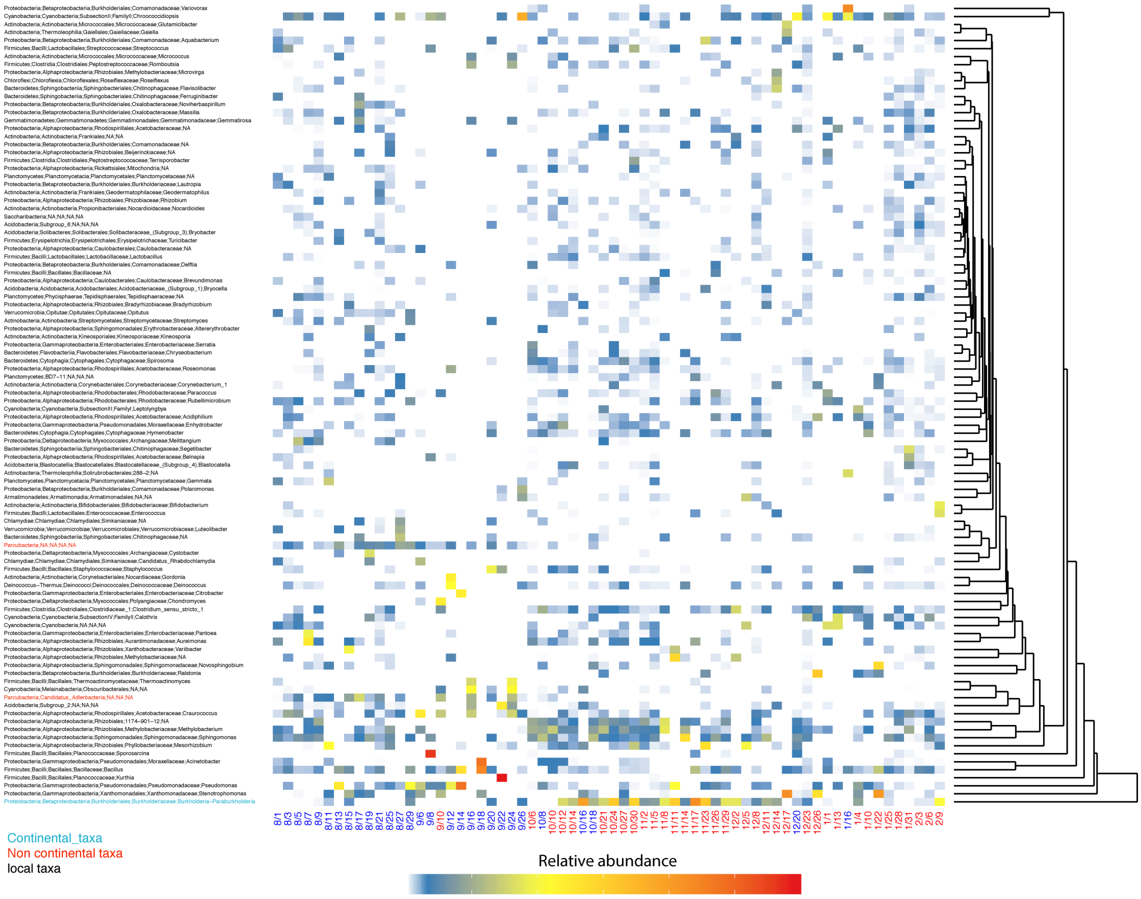


SI Fig. 2: Seasonal change of bacteria at the genus level.

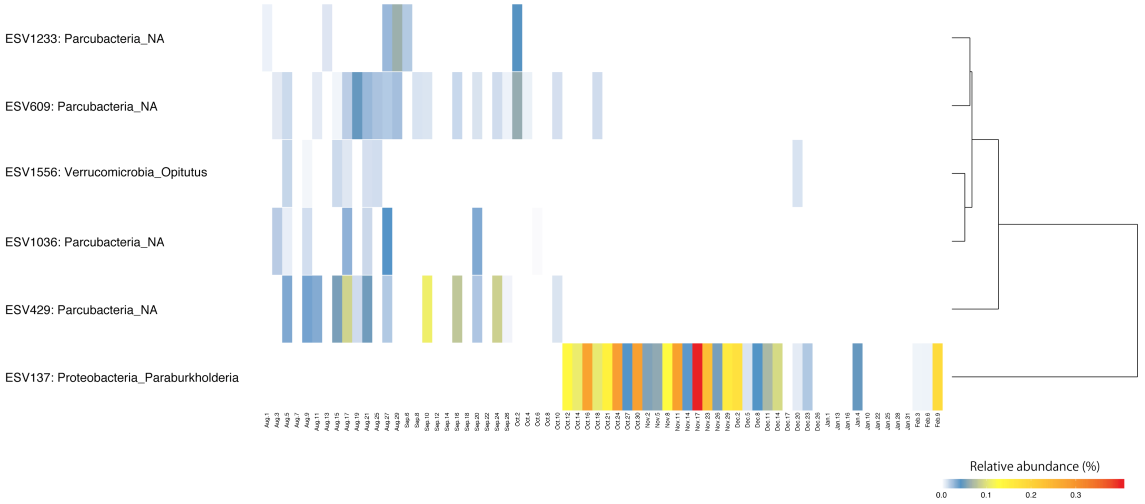


SI Fig. 3: Seasonal change of 6 ASVs, which were strongly affected by air mass change in early October.

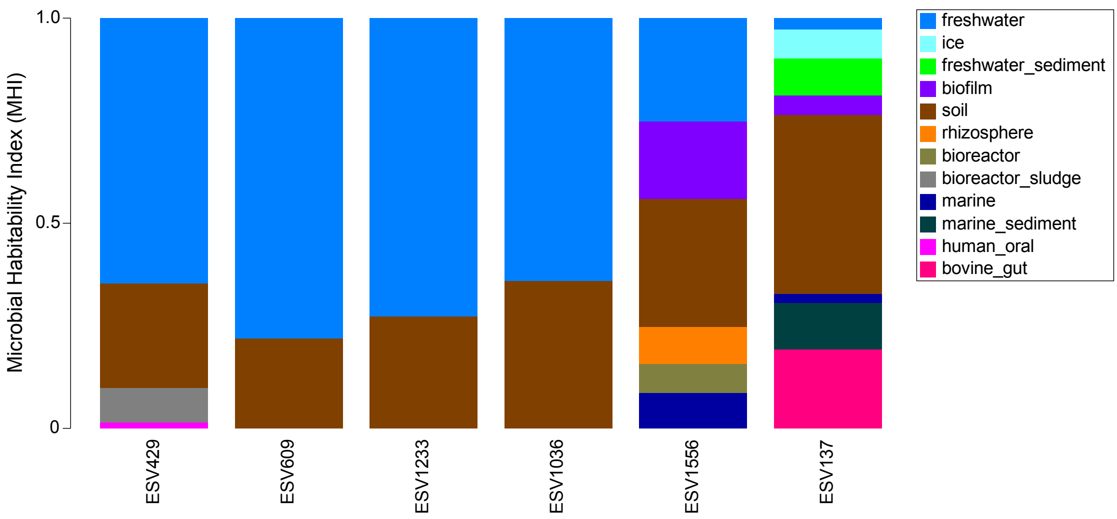


SI Fig. 4: Microbial habitability index for 6 ASVs estimated by metametaDB, which infers the possibility to find microbes in each environment.

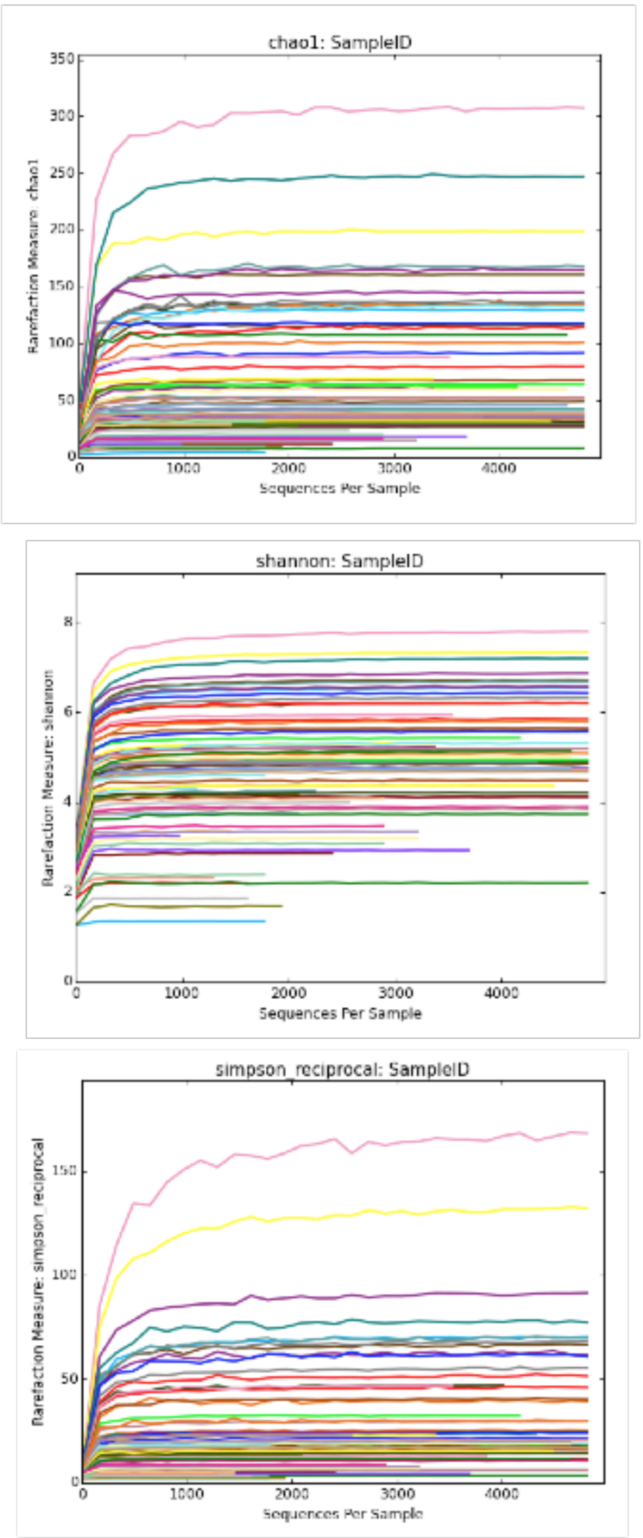


SI Fig. 5: Rarefaction curves of three alpha diversity methods.

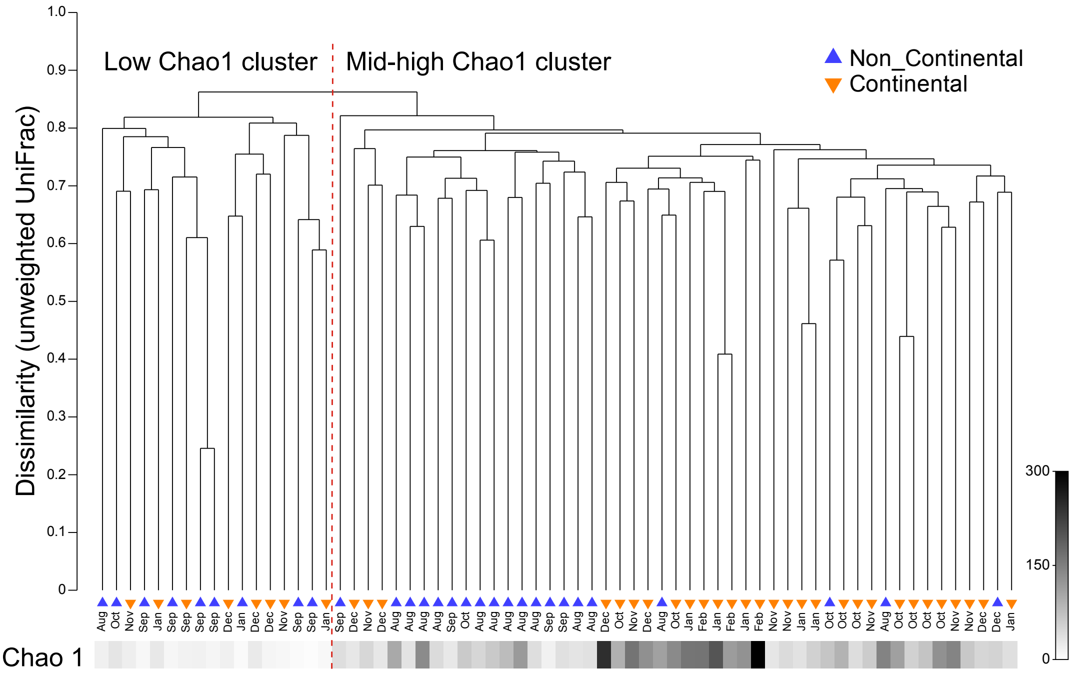


SI Fig. 6: Unweighted Pair Group Method with Arithmetic Mean (UPGMA) dendrogram of dissimilarity by unweighted-UniFrac, shown with changes of Chao1 (gray bar in the bottom).

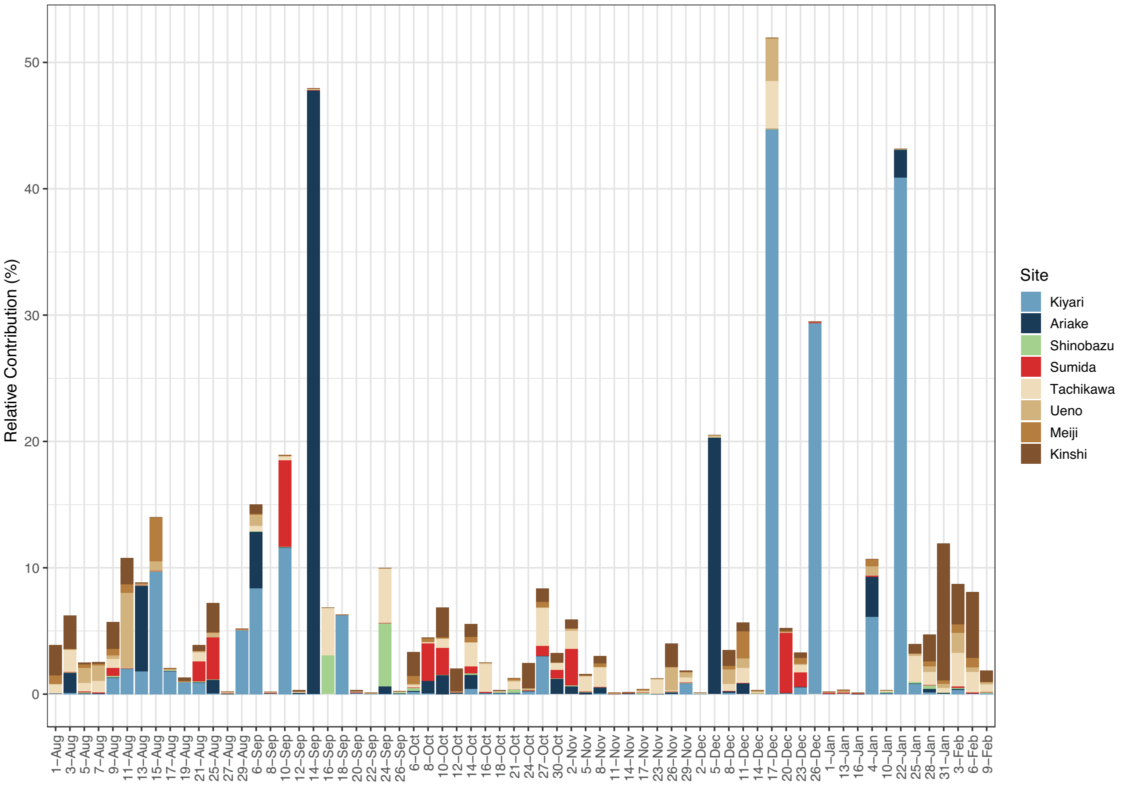


SI Fig. 7: Seasonal change of estimated contribution from 8 potential sites in 4 categories (bay, soil, pond and river) by source tracking analysis.

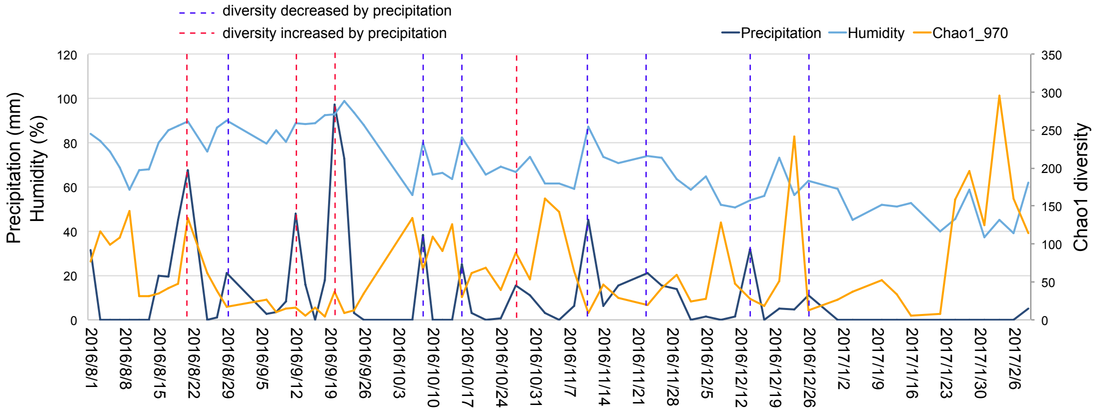


SI Fig.8: Changes of Chao 1 index and potentially related meteorology factors (precipitation and humidity). Blue broken lines show sampling periods when Chao 1 diversity decreased during precipitation in normal rain events. Red broken lines show sampling periods when Chao 1 diversity increased during precipitation, primarily occurring during a historically heavy rain event.
