## Supplemental Table 1 for "Seasonal changes of airborne bacterial communities over Tokyo and influence of local meteorology"

|  | Site name | Sample category | Latitude | Longtitude | Distance from Tokyo Skytree (km) | Aspect from Tokyo Skytree |
| --- | --- | --- | --- | --- | --- | --- |
| DNA sample | Tokyo Skytree | Air | 35.71013889 | 139.8108333 | 0 | - |
|  | Ariake | Bay | 35.628475 | 139.7862472 | 9.3 | SW |
|  | Kiyari | Bay | 35.64550556 | 139.7915306 | 7.4 | SSW |
|  | Kinshi | Soil | 35.69857778 | 139.8155917 | 1.3 | S |
|  | Ueno | Soil | 35.71449444 | 139.7732444 | 3.4 | W |
|  | Meiji | Soil | 35.67115556 | 139.7014278 | 10.9 | SW |
|  | Tachikawa | Soil | 35.71243611 | 139.4072833 | 36.6 | W |
|  | Sumida | River | 35.70837222 | 139.7967028 | 1.3 | W |
|  | Shinobazu | Pond | 35.71246111 | 139.7704278 | 3.7 | W |
| Environmental data | Edogawa rinkai | Meteorogical station | 35.63888333 | 139.8636278 | 9.3 | SE |
|  | Tokyo | Meteorogical station | 35.69179722 | 139.7513361 | 5.7 | SWW |
|  | Tokyo Bay | Ocean observatory | 35.57774722 | 139.8174278 | 14.5 | S |

SI Table 1: Information of sampling sites and meteorological stations
