## Supplemental Table2 for "Seasonal changes of airborne bacterial communities over Tokyo and influence of local meteorology"

|  | Sampling |  |  |  |  | Sample category |  |  | Meteorological data |  |  |  |  |  | Possible contribution by Sourcetracking analysis (%) | | | |  |  |  |  |  |  |  |  |  | Aipha diversity |  |  |
| --- | --- | --- | --- | --- | --- | --- | --- | --- | --- | --- | --- | --- | --- | --- | --- | --- | --- | --- | --- | --- | --- | --- | --- | --- | --- | --- | --- | --- | --- | --- |
|  | Start date | End date | Sampling time (h) | Sampling volume (m3) | Average sampling volume（L/min） | HYSPLIT | Month | Number of sequences | Precipitation (mm) | Air temp (℃) | Wind speed (m/s) | Sunlight duration (h) | Humidity (mm) | Wave_height (m) | Ariake | Kinshi | Kiyari | Meiji | Tachikawa | Shinobazu | Sumida | Ueno | **soil** | **bay** | **river** | **pond** | **total** | Chao1 | Shannon | Simpson_reciprocal |
| 05W1 | 8/1/16 | 8/2/16 | 48 | 86.398 | 30.0 | Non Continental | Aug | 14889 | 31.5 | 26.8 | 3.0 | 2.7 | 84.0 | 0.2 | 0.0 | 2.4 | 0.0 | 0.7 | 0.7 | 0.0 | 0.0 | 0.1 | 3.8 | 0.0 | 0.0 | 0.0 | 3.9 | 77.2 | 5.8 | 44.6 |
| 05W2 | 8/3/16 | 8/4/16 | 48 | 85.037 | 29.5 | Non Continental | Aug | 9279 | 0.0 | 27.9 | 3.6 | 9.0 | 81.0 | 0.2 | 1.6 | 2.6 | 0.1 | 0.0 | 1.8 | 0.0 | 0.0 | 0.0 | 4.5 | 1.7 | 0.0 | 0.0 | 6.2 | 117.1 | 6.3 | 58.4 |
| 05W3 | 8/5/16 | 8/6/16 | 48 | 85.771 | 29.8 | Non Continental | Aug | 12235 | 0.0 | 28.3 | 4.9 | 12.3 | 76.0 | 0.2 | 0.1 | 0.1 | 0.0 | 0.3 | 0.7 | 0.1 | 0.1 | 1.2 | 2.3 | 0.1 | 0.1 | 0.1 | 2.5 | 99.1 | 5.8 | 30.1 |
| 05W4 | 8/7/16 | 8/8/16 | 48 | 86.051 | 29.9 | Non Continental | Aug | 4764 | 0.0 | 28.8 | 4.1 | 8.1 | 69.0 | 0.2 | 0.0 | 0.1 | 0.0 | 0.1 | 0.9 | 0.0 | 0.1 | 1.3 | 2.4 | 0.0 | 0.1 | 0.0 | 2.6 | 108.4 | 5.0 | 11.6 |
| 05W5 | 8/9/16 | 8/10/16 | 48 | 78.532 | 27.3 | Non Continental | Aug | 11582 | 0.0 | 30.2 | 3.4 | 6.2 | 59.0 | 0.2 | 0.0 | 2.1 | 1.3 | 0.5 | 0.7 | 0.1 | 0.7 | 0.3 | 3.6 | 1.4 | 0.7 | 0.1 | 5.7 | 143.1 | 6.5 | 60.4 |
| 05W6 | 8/11/16 | 8/12/16 | 30 | 44.654 | 24.8 | Non Continental | Aug | 4530 | 0.0 | 26.8 | 3.2 | 5.1 | 67.5 | 0.2 | 0.0 | 2.1 | 2.0 | 0.6 | 0.0 | 0.0 | 0.0 | 5.9 | 8.7 | 2.0 | 0.0 | 0.0 | 10.8 | 31.5 | 4.4 | 15.3 |
| 05W7 | 8/13/16 | 8/14/16 | 48 | 75.381 | 26.2 | Non Continental | Aug | 1882 | 0.0 | 26.3 | 3.4 | 7.7 | 68.0 | 0.2 | 6.8 | 0.1 | 1.8 | 0.0 | 0.0 | 0.0 | 0.1 | 0.0 | 0.1 | 8.6 | 0.1 | 0.0 | 8.8 | 31.8 | 4.6 | 17.7 |
| 06W1 | 8/15/16 | 8/16/16 | 48 | 86.398 | 30.0 | Non Continental | Aug | 3888 | 20.0 | 27.1 | 4.5 | 2.2 | 80.0 | 0.2 | 0.0 | 0.0 | 9.7 | 3.5 | 0.0 | 0.0 | 0.0 | 0.7 | 4.3 | 9.7 | 0.0 | 0.0 | 14.0 | 34.8 | 4.4 | 12.9 |
| 06W2 | 8/17/16 | 8/18/16 | 1.02 | 1.874 | 30.6 | Non Continental | Aug | 4361 | 19.5 | 27.6 | 5.8 | 5.8 | 85.5 | 0.3 | 0.0 | 0.0 | 1.8 | 0.1 | 0.0 | 0.0 | 0.0 | 0.1 | 0.2 | 1.8 | 0.0 | 0.0 | 2.1 | 41.9 | 4.9 | 23.4 |
| 06W3 | 8/19/16 | 8/20/16 | 48 | 84.189 | 29.2 | Non Continental | Aug | 7516 | 45.0 | 27.2 | 3.8 | 5.2 | 87.5 | 0.2 | 0.0 | 0.2 | 0.9 | 0.0 | 0.0 | 0.1 | 0.0 | 0.0 | 0.3 | 0.9 | 0.0 | 0.1 | 1.3 | 47.5 | 4.8 | 17.8 |
| 06W4 | 8/21/16 | 8/22/16 | 48 | 86.07 | 29.9 | Non Continental | Aug | 7290 | 67.5 | 27.4 | 5.5 | 4.2 | 89.5 | 0.4 | 0.0 | 0.5 | 0.9 | 0.0 | 0.7 | 0.1 | 1.6 | 0.1 | 1.3 | 0.9 | 1.6 | 0.1 | 3.9 | 134.0 | 6.5 | 65.4 |
| 07W1 | 8/25/16 | 8/26/16 | 48 | 85.965 | 29.8 | Non Continental | Aug | 4319 | 0.0 | 28.0 | 3.2 | 12.5 | 76.0 | 0.2 | 1.1 | 2.3 | 0.1 | 0.0 | 0.0 | 0.0 | 3.3 | 0.3 | 2.7 | 1.1 | 3.3 | 0.0 | 7.2 | 62.1 | 5.4 | 32.0 |
| 07W2 | 8/27/16 | 8/28/16 | 48 | 86.398 | 30.0 | Non Continental | Aug | 5044 | 1.0 | 23.7 | 4.1 | 0.8 | 87.0 | 0.3 | 0.0 | 0.0 | 0.0 | 0.0 | 0.1 | 0.1 | 0.0 | 0.0 | 0.1 | 0.0 | 0.0 | 0.1 | 0.2 | 37.8 | 4.7 | 20.2 |
| 07W3 | 8/29/16 | 8/30/16 | 35 | 55.228 | 26.3 | Non Continental | Aug | 2225 | 21.0 | 26.1 | 5.3 | 2.1 | 90.0 | 0.2 | 0.0 | 0.0 | 5.1 | 0.0 | 0.0 | 0.0 | 0.0 | 0.0 | 0.1 | 5.1 | 0.0 | 0.0 | 5.2 | 17.0 | 3.7 | 10.8 |
| 07W7 | 9/6/16 | 9/7/16 | 48.00 | 71.999 | 25.0 | Non Continental | Sep | 2938 | 2.5 | 27.3 | 6.5 | 6.8 | 79.5 | 0.3 | 4.4 | 0.7 | 8.4 | 0.1 | 0.4 | 0.0 | 0.0 | 0.9 | 2.1 | 12.8 | 0.0 | 0.0 | 15.0 | 27.0 | 4.0 | 13.2 |
| 08W1 | 9/8/16 | 9/9/16 | 48.00 | 71.998 | 25.0 | Non Continental | Sep | 1808 | 3.5 | 27.4 | 4.0 | 6.9 | 85.5 | 0.2 | 0.0 | 0.0 | 0.0 | 0.0 | 0.0 | 0.0 | 0.0 | 0.0 | 0.1 | 0.1 | 0.0 | 0.0 | 0.2 | 10.0 | 2.4 | 3.2 |
| 08W2 | 9/10/16 | 9/11/16 | 48.00 | 71.999 | 25.0 | Continental | Sep | 3293 | 8.5 | 24.8 | 3.0 | 3.2 | 80.5 | 0.2 | 0.0 | 0.0 | 11.6 | 0.1 | 0.3 | 0.0 | 6.8 | 0.0 | 0.4 | 11.6 | 6.8 | 0.0 | 18.9 | 15.0 | 3.3 | 8.0 |
| 08W3 | 9/12/16 | 9/13/16 | 48.00 | 71.995 | 25.0 | Non Continental | Sep | 3228 | 48.0 | 23.9 | 2.6 | 0.7 | 89.0 | 0.2 | 0.1 | 0.1 | 0.0 | 0.1 | 0.1 | 0.0 | 0.0 | 0.0 | 0.2 | 0.1 | 0.0 | 0.0 | 0.3 | 16.0 | 3.2 | 6.8 |
| 08W4 | 9/14/16 | 9/15/16 | 32.20 | 42.084 | 21.8 | Non Continental | Sep | 1685 | 16.0 | 24.0 | 1.9 | 0.3 | 88.5 | 0.2 | 47.8 | 0.0 | 0.0 | 0.0 | 0.0 | 0.0 | 0.0 | 0.0 | 0.1 | 47.8 | 0.0 | 0.0 | 48.0 | 6.0 | 1.9 | 3.2 |
| 08W5 | 9/16/16 | 9/17/16 | 48 | 64.754 | 22.5 | Non Continental | Sep | 2931 | 0.0 | 24.4 | 2.3 | 3.8 | 89.0 | 0.2 | 0.0 | 0.0 | 0.0 | 0.0 | 3.7 | 3.0 | 0.0 | 0.0 | 3.8 | 0.0 | 0.0 | 3.0 | 6.9 | 16.0 | 3.5 | 8.9 |
| 09W1 | 9/18/16 | 9/19/16 | 14.03 | 17.253 | 20.5 | Non Continental | Sep | 1784 | 18.0 | 23.3 | 3.8 | 0.0 | 92.5 | 0.2 | 0.0 | 0.0 | 6.3 | 0.0 | 0.0 | 0.0 | 0.0 | 0.0 | 0.0 | 6.3 | 0.0 | 0.0 | 6.3 | 4.0 | 1.3 | 2.3 |
| 09W2 | 9/20/16 | 9/21/16 | 48.00 | 71.999 | 25.0 | Non Continental | Sep | 8451 |  | 20.0 | 3.6 | 0.1 | 93.0 | 0.2 | 0.0 | 0.1 | 0.0 | 0.0 | 0.1 | 0.0 | 0.0 | 0.0 | 0.2 | 0.1 | 0.0 | 0.0 | 0.3 | 37.6 | 4.6 | 16.9 |
| 09W3 | 9/22/16 | 9/23/16 | 48.00 | 71.998 | 25.0 | Non Continental | Sep | 1980 | 73.0 | 20.4 | 2.2 | 0.0 | 99.0 | 0.2 | 0.0 | 0.0 | 0.0 | 0.0 | 0.0 | 0.0 | 0.0 | 0.0 | 0.1 | 0.0 | 0.0 | 0.0 | 0.1 | 8.9 | 1.7 | 2.4 |
| 09W4 | 9/24/16 | 9/25/16 | 48.00 | 71.999 | 25.0 | Non Continental | Sep | 1068 | 3.0 | 22.8 | 1.7 | 2.3 | 93.5 | 0.2 | 0.6 | 0.0 | 0.0 | 0.0 | 4.3 | 5.0 | 0.0 | 0.0 | 4.3 | 0.6 | 0.0 | 5.0 | 10.0 | 12.0 | 3.3 | 8.2 |
| 09W5 | 9/26/16 | 9/27/16 | 48.00 | 71.999 | 25.0 | Non Continental | Sep | 10655 | 0.0 | 25.4 | 2.1 | 5.1 | 88.0 | 0.2 | 0.0 | 0.0 | 0.0 | 0.0 | 0.0 | 0.1 | 0.0 | 0.0 | 0.1 | 0.1 | 0.0 | 0.1 | 0.3 | 35.3 | 3.8 | 6.5 |
| 10W3 | 10/2/16 | 10/3/16 | 48.00 | 86.398 | 30.0 | Non Continental | Oct | 8865 | 5.5 | 22.9 | 2.8 | 3.8 | 86.0 |  | 0.4 | 0.0 | 0.0 | 0.0 | 0.0 | 0.0 | 1.4 | 0.0 | 0.1 | 0.4 | 1.4 | 0.0 | 1.9 | 41.8 | 4.9 | 24.1 |
| 10W4 | 10/4/16 | 10/5/16 | 48.00 | 76.477 | 26.6 | Continental | Oct | 10658 | 1.0 | 24.5 | 3.6 | 4.2 | 78.5 |  | 0.0 | 0.0 | 0.0 | 0.0 | 5.3 | 0.0 | 0.0 | 0.1 | 5.4 | 0.0 | 0.0 | 0.0 | 5.5 | 31.0 | 4.1 | 11.1 |
| 10W5 | 10/6/16 | 10/7/16 | 48.00 | 85.331 | 29.6 | Continental | Oct | 4102 | 0.0 | 24.1 | 4.0 | 7.6 | 56.5 | 0.3 | 0.1 | 1.9 | 0.2 | 0.6 | 0.2 | 0.2 | 0.0 | 0.1 | 2.8 | 0.3 | 0.0 | 0.2 | 3.3 | 134.6 | 6.1 | 45.2 |
| 11W1 | 10/8/16 | 10/9/16 | 48.00 | 43.199 | 15.0 | Non Continental | Oct | 14256 | 38.5 | 22.3 | 5.3 | 0.4 | 80.0 | 0.3 | 1.0 | 0.1 | 0.1 | 0.3 | 0.1 | 0.0 | 2.9 | 0.0 | 0.5 | 1.1 | 2.9 | 0.0 | 4.5 | 66.8 | 5.6 | 39.5 |
| 11W2 | 10/10/16 | 10/11/16 | 48.00 | 43.199 | 15.0 | Continental | Oct | 19913 | 0.0 | 18.2 | 3.1 | 0.0 | 65.5 | 0.2 | 1.5 | 2.4 | 0.0 | 0.1 | 0.7 | 0.0 | 2.2 | 0.0 | 3.2 | 1.5 | 2.2 | 0.0 | 6.9 | 109.4 | 6.1 | 49.3 |
| 11W3 | 10/12/16 | 10/13/16 | 48.00 | 43.199 | 15.0 | Continental | Oct | 11251 | 0.0 | 17.6 | 2.8 | 4.3 | 66.5 | 0.3 | 0.1 | 1.8 | 0.0 | 0.0 | 0.0 | 0.0 | 0.0 | 0.0 | 1.9 | 0.1 | 0.0 | 0.0 | 2.0 | 91.0 | 5.5 | 24.0 |
| 11W4 | 10/14/16 | 10/15/16 | 48.00 | 43.199 | 15.0 | Continental | Oct | 9415 | 0.0 | 17.1 | 2.5 | 7.7 | 63.5 | 0.2 | 1.1 | 1.0 | 0.4 | 0.4 | 1.9 | 0.1 | 0.6 | 0.0 | 3.3 | 1.5 | 0.6 | 0.1 | 5.5 | 126.4 | 6.1 | 40.1 |
| 11W5 | 10/16/16 | 10/17/16 | 48.00 | 43.199 | 15.0 | Non Continental | Oct | 5534 | 25.0 | 18.6 | 2.3 | 3.4 | 82.5 | 0.2 | 0.0 | 0.0 | 0.1 | 0.0 | 2.2 | 0.0 | 0.1 | 0.1 | 2.3 | 0.1 | 0.1 | 0.0 | 2.5 | 30.0 | 3.7 | 6.1 |
| 12W1 | 10/18/16 | 10/20/16 | 72.00 | 64.799 | 15.0 | Non Continental | Oct | 3442 | 3.0 | 21.6 | 2.5 | 6.3 | 75.7 | 0.2 | 0.0 | 0.1 | 0.1 | 0.0 | 0.0 | 0.1 | 0.0 | 0.1 | 0.2 | 0.1 | 0.0 | 0.1 | 0.3 | 61.2 | 5.2 | 23.5 |
| 12W2 | 10/21/16 | 10/23/16 | 72.00 | 64.799 | 15.0 | Continental | Oct | 3512 | 0.0 | 18.0 | 3.0 | 3.6 | 65.7 | 0.2 | 0.0 | 0.0 | 0.1 | 0.1 | 0.6 | 0.3 | 0.0 | 0.1 | 0.9 | 0.1 | 0.0 | 0.3 | 1.3 | 68.5 | 5.3 | 22.9 |
| 12W3 | 10/24/16 | 10/26/16 | 60.00 | 53.999 | 15.0 | Continental | Oct | 1266 | 0.5 | 16.6 | 2.5 | 8.0 | 69.3 | 0.2 | 0.0 | 2.0 | 0.2 | 0.0 | 0.0 | 0.0 | 0.0 | 0.1 | 2.2 | 0.2 | 0.0 | 0.0 | 2.5 | 38.9 | 4.3 | 8.9 |
| 13W1 | 10/27/16 | 10/29/16 | 72.00 | 53.999 | 12.5 | Continental | Oct | 3597 | 15.5 | 16.1 | 3.0 | 2.8 | 66.7 | 0.2 | 0.0 | 1.1 | 3.0 | 0.4 | 3.0 | 0.0 | 0.8 | 0.0 | 4.5 | 3.0 | 0.8 | 0.0 | 8.4 | 88.1 | 5.9 | 46.3 |
| 13W2 | 10/30/16 | 11/1/16 | 72.00 | 64.799 | 15.0 | Continental | Oct | 2372 | 11.0 | 13.5 | 2.4 | 2.5 | 73.7 | 0.2 | 1.2 | 0.7 | 0.0 | 0.0 | 0.5 | 0.1 | 0.7 | 0.0 | 1.3 | 1.2 | 0.7 | 0.1 | 3.2 | 53.1 | 4.2 | 8.7 |
| 13W3 | 11/2/16 | 11/4/16 | 72.00 | 64.799 | 15.0 | Continental | Nov | 5406 | 3.0 | 13.2 | 3.0 | 6.5 | 61.7 | 0.2 | 0.6 | 0.7 | 0.0 | 0.0 | 1.4 | 0.1 | 2.9 | 0.2 | 2.3 | 0.6 | 2.9 | 0.1 | 5.9 | 160.4 | 6.6 | 61.9 |
| 13W4 | 11/5/16 | 11/7/16 | 72.00 | 64.799 | 15.0 | Continental | Nov | 8407 | 0.0 | 14.0 | 3.5 | 8.6 | 61.7 | 0.2 | 0.1 | 0.1 | 0.0 | 0.1 | 1.2 | 0.0 | 0.1 | 0.0 | 1.3 | 0.1 | 0.1 | 0.0 | 1.6 | 142.1 | 6.2 | 53.0 |
| 13W5 | 11/8/16 | 11/10/16 | 72.00 | 64.799 | 15.0 | Continental | Nov | 4964 | 6.5 | 10.4 | 3.7 | 2.6 | 59.3 | 0.2 | 0.5 | 0.6 | 0.1 | 0.3 | 1.5 | 0.0 | 0.0 | 0.0 | 2.4 | 0.5 | 0.0 | 0.0 | 3.0 | 63.5 | 4.9 | 17.9 |
| 14W1 | 11/11/16 | 11/13/16 | 74.00 | 66.599 | 15.0 | Continental | Nov | 1318 | 45.0 | 12.6 | 2.7 | 4.7 | 87.3 | 0.2 | 0.0 | 0.0 | 0.0 | 0.0 | 0.0 | 0.0 | 0.0 | 0.0 | 0.1 | 0.0 | 0.0 | 0.0 | 0.1 | 9.0 | 2.3 | 4.3 |
| 14W2 | 11/14/16 | 11/16/16 | 72.00 | 64.799 | 15.0 | Continental | Nov | 4773 | 6.5 | 14.3 | 2.4 | 0.5 | 73.7 | 0.2 | 0.0 | 0.0 | 0.0 | 0.0 | 0.0 | 0.0 | 0.0 | 0.0 | 0.1 | 0.0 | 0.0 | 0.0 | 0.2 | 46.1 | 4.7 | 19.1 |
| 14W3 | 11/17/16 | 11/19/16 | 72.00 | 66.301 | 15.3 | Continental | Nov | 1572 | 15.5 | 11.5 | 2.2 | 5.7 | 71.0 | 0.2 | 0.0 | 0.0 | 0.0 | 0.1 | 0.1 | 0.1 | 0.0 | 0.1 | 0.3 | 0.0 | 0.0 | 0.1 | 0.4 | 29.0 | 3.3 | 4.8 |
| 14W5 | 11/23/16 | 11/25/16 | 72.00 | 64.799 | 15.0 | Continental | Nov | 3046 | 21.0 | 6.7 | 2.8 | 3.2 | 74.0 | 0.2 | 0.0 | 0.0 | 0.0 | 0.0 | 1.1 | 0.0 | 0.0 | 0.0 | 1.2 | 0.0 | 0.0 | 0.0 | 1.2 | 19.8 | 3.1 | 6.0 |
| 14W6 | 11/26/16 | 11/28/16 | 72.00 | 64.8 | 15.0 | Continental | Nov | 2656 | 15.5 | 10.4 | 3.0 | 3.1 | 73.3 | 0.2 | 0.1 | 1.9 | 0.0 | 0.0 | 0.1 | 0.1 | 0.0 | 1.9 | 3.8 | 0.1 | 0.0 | 0.1 | 4.0 | 42.0 | 4.9 | 22.9 |
| 14W7 | 11/29/16 | 12/1/16 | 153.54 | 138.519 | 15.0 | Continental | Nov | 4674 | 14.0 | 10.1 | 3.4 | 4.8 | 63.7 | 0.2 | 0.0 | 0.1 | 0.9 | 0.0 | 0.4 | 0.0 | 0.0 | 0.4 | 0.9 | 0.9 | 0.0 | 0.0 | 1.8 | 58.9 | 4.9 | 18.2 |
| 15W1 | 12/2/16 | 12/4/16 | 72.00 | 64.799 | 15.0 | Continental | Dec | 2577 | 0.0 | 12.1 | 2.4 | 7.6 | 59.0 | 0.2 | 0.0 | 0.0 | 0.0 | 0.0 | 0.0 | 0.0 | 0.0 | 0.0 | 0.1 | 0.0 | 0.0 | 0.0 | 0.1 | 23.9 | 4.0 | 12.2 |
| 15W2 | 12/5/16 | 12/7/16 | 72.00 | 64.799 | 15.0 | Continental | Dec | 6037 | 1.5 | 10.8 | 2.9 | 6.2 | 65.0 | 0.2 | 20.3 | 0.0 | 0.0 | 0.0 | 0.0 | 0.0 | 0.0 | 0.1 | 0.2 | 20.3 | 0.0 | 0.0 | 20.5 | 28.0 | 3.9 | 11.0 |
| 15W3 | 12/8/16 | 12/10/16 | 72.00 | 64.798 | 15.0 | Continental | Dec | 6246 | 0.0 | 10.1 | 4.2 | 9.2 | 52.0 | 0.2 | 0.1 | 1.2 | 0.1 | 0.3 | 0.6 | 0.0 | 0.0 | 1.1 | 3.2 | 0.2 | 0.0 | 0.0 | 3.5 | 127.9 | 6.5 | 66.4 |
| 15W4 | 12/11/16 | 12/13/16 | 72.00 | 65.202 | 15.1 | Continental | Dec | 2564 | 1.5 | 8.2 | 2.7 | 6.2 | 51.0 | 0.2 | 0.8 | 0.7 | 0.0 | 2.1 | 1.2 | 0.0 | 0.0 | 0.8 | 4.8 | 0.8 | 0.0 | 0.0 | 5.7 | 47.1 | 4.9 | 23.2 |
| 15W5 | 12/14/16 | 12/16/16 | 72.00 | 64.799 | 15.0 | Continental | Dec | 2108 | 32.0 | 6.6 | 3.9 | 4.3 | 54.0 | 0.2 | 0.0 | 0.0 | 0.0 | 0.1 | 0.1 | 0.1 | 0.0 | 0.1 | 0.2 | 0.0 | 0.0 | 0.1 | 0.3 | 28.0 | 4.2 | 13.6 |
| 15W6 | 12/17/16 | 12/19/16 | 72.00 | 64.799 | 15.0 | Continental | Dec | 3713 | 0.0 | 9.1 | 1.7 | 8.9 | 56.0 | 0.2 | 0.0 | 0.0 | 44.7 | 0.0 | 3.8 | 0.1 | 0.0 | 3.4 | 7.2 | 44.7 | 0.0 | 0.1 | 52.0 | 17.8 | 2.9 | 4.6 |
| 15W7 | 12/20/16 | 12/22/16 | 72.00 | 64.713 | 15.0 | Non Continental | Dec | 5440 | 5.0 | 11.4 | 3.2 | 4.6 | 73.3 | 0.2 | 0.0 | 0.2 | 0.0 | 0.1 | 0.1 | 0.0 | 4.7 | 0.0 | 0.4 | 0.1 | 4.7 | 0.0 | 5.2 | 51.0 | 5.2 | 29.0 |
| 16W1 | 12/23/16 | 12/25/16 | 72.00 | 64.799 | 15.0 | Continental | Dec | 7929 | 4.5 | 10.2 | 3.7 | 5.9 | 56.3 | 0.2 | 0.0 | 0.4 | 0.5 | 0.5 | 0.6 | 0.0 | 1.1 | 0.1 | 1.6 | 0.6 | 1.1 | 0.0 | 3.3 | 241.5 | 7.0 | 75.2 |
| 16W2 | 12/26/16 | 12/28/16 | 72.00 | 64.799 | 15.0 | Continental | Dec | 2495 | 11.0 | 8.3 | 4.2 | 3.5 | 63.0 | 0.3 | 0.0 | 0.0 | 29.3 | 0.0 | 0.0 | 0.0 | 0.1 | 0.0 | 0.1 | 29.4 | 0.1 | 0.0 | 29.5 | 12.0 | 2.8 | 5.3 |
| 16W4 | 1/1/17 | 1/3/17 | 72.00 | 64.799 | 15.0 | Continental | Jan | 4872 | 0.0 | 8.0 | 2.1 | 8.4 | 59.3 | 0.1 | 0.0 | 0.0 | 0.0 | 0.0 | 0.0 | 0.0 | 0.1 | 0.0 | 0.1 | 0.0 | 0.1 | 0.0 | 0.2 | 26.3 | 4.2 | 14.2 |
| 17W1 | 1/13/17 | 1/15/17 | 72.00 | 65.572 | 15.2 | Continental | Jan | 6044 | 0.0 | 6.7 | 3.9 | 9.2 | 45.0 | 0.2 | 0.0 | 0.0 | 0.0 | 0.1 | 0.0 | 0.0 | 0.1 | 0.1 | 0.2 | 0.1 | 0.1 | 0.0 | 0.3 | 33.6 | 4.4 | 15.6 |
| 17W2 | 1/16/17 | 1/18/17 | 72.00 | 64.799 | 15.0 | Non Continental | Jan | 1653 | 0.0 | 7.9 | 2.9 | 9.0 | 52.0 | 0.2 | 0.0 | 0.0 | 0.0 | 0.0 | 0.0 | 0.0 | 0.0 | 0.0 | 0.1 | 0.0 | 0.0 | 0.0 | 0.1 | 6.0 | 2.2 | 3.8 |
| 17W3 | 1/4/17 | 1/6/17 | 72.00 | 64.799 | 15.0 | Continental | Jan | 7175 | 0.0 | 3.8 | 3.0 | 6.8 | 51.3 | 0.2 | 3.2 | 0.0 | 6.1 | 0.5 | 0.0 | 0.0 | 0.1 | 0.7 | 1.3 | 9.3 | 0.1 | 0.0 | 10.7 | 37.1 | 4.7 | 21.5 |
| 17W5 | 1/10/17 | 1/12/17 | 72.00 | 64.799 | 15.0 | Continental | Jan | 9022 | 0.0 | 5.6 | 3.1 | 9.4 | 52.7 | 0.2 | 0.0 | 0.0 | 0.1 | 0.0 | 0.1 | 0.1 | 0.0 | 0.0 | 0.1 | 0.1 | 0.0 | 0.1 | 0.3 | 52.6 | 5.1 | 24.8 |
| 19W1 | 1/22/17 | 1/24/17 | 72.00 | 64.8 | 15.0 | Continental | Jan | 4944 | 0.0 | 4.7 | 3.4 | 8.6 | 40.0 | 0.2 | 2.2 | 0.0 | 40.9 | 0.0 | 0.0 | 0.0 | 0.0 | 0.0 | 0.0 | 43.1 | 0.0 | 0.0 | 43.2 | 7.7 | 2.2 | 3.6 |
| 19W2 | 1/25/17 | 1/27/17 | 72.00 | 64.799 | 15.0 | Continental | Jan | 9978 | 0.0 | 6.9 | 4.7 | 8.1 | 45.7 | 0.3 | 0.0 | 0.7 | 0.8 | 0.0 | 2.1 | 0.1 | 0.0 | 0.2 | 3.0 | 0.9 | 0.0 | 0.1 | 4.0 | 159.3 | 6.8 | 85.2 |
| 19W3 | 1/28/17 | 1/30/17 | 72.00 | 64.799 | 15.0 | Continental | Jan | 6478 | 0.0 | 9.6 | 2.8 | 7.4 | 58.7 | 0.2 | 0.3 | 2.1 | 0.1 | 0.4 | 1.0 | 0.3 | 0.0 | 0.4 | 4.0 | 0.4 | 0.0 | 0.3 | 4.7 | 195.8 | 7.2 | 120.1 |
| 19W4 | 1/31/17 | 2/2/17 | 72.00 | 64.799 | 15.0 | Continental | Jan | 7697 | 0.0 | 5.9 | 3.9 | 9.2 | 37.3 | 0.2 | 0.1 | 10.8 | 0.0 | 0.3 | 0.3 | 0.0 | 0.0 | 0.3 | 11.8 | 0.1 | 0.0 | 0.0 | 11.9 | 125.4 | 5.2 | 18.6 |
| 19W5 | 2/3/17 | 2/5/17 | 72.00 | 64.799 | 15.0 | Continental | Feb | 15482 | 0.0 | 7.8 | 3.1 | 7.3 | 45.3 | 0.2 | 0.1 | 3.2 | 0.4 | 0.7 | 2.7 | 0.1 | 0.1 | 1.6 | 8.1 | 0.4 | 0.1 | 0.1 | 8.7 | 295.4 | 7.6 | 150.8 |
| 20W1 | 2/6/17 | 2/8/17 | 72.00 | 64.799 | 15.0 | Continental | Feb | 11213 | 0.0 | 7.3 | 4.3 | 9.2 | 39.0 | 0.2 | 0.0 | 5.2 | 0.0 | 0.8 | 1.6 | 0.0 | 0.1 | 0.4 | 8.0 | 0.1 | 0.1 | 0.0 | 8.1 | 160.2 | 6.6 | 66.4 |
| 20W2 | 2/9/17 | 2/11/17 | 72.00 | 64.799 | 15.0 | Continental | Feb | 6343 | 5.0 | 4.0 | 3.3 | 4.4 | 62.0 | 0.2 | 0.0 | 0.9 | 0.1 | 0.0 | 0.6 | 0.1 | 0.0 | 0.1 | 1.7 | 0.1 | 0.0 | 0.1 | 1.9 | 114.2 | 4.8 | 11.4 |
| Minimum |  |  |  |  |  |  |  | 1068.0 | 0.0 | 3.8 | 1.7 | 0.0 | 37.3 | 0.1 | 0.0 | 0.0 | 0.0 | 0.0 | 0.0 | 0.0 | 0.0 | 0.0 | 0.0 | 0.0 | 0.0 | 0.0 | 0.1 | 4.0 | 1.3 | 2.3 |
| Maximum |  |  |  |  |  |  |  | 19913.0 | 97.5 | 30.2 | 6.5 | 12.5 | 99.0 | 0.4 | 47.8 | 10.8 | 44.7 | 3.5 | 5.3 | 5.0 | 6.8 | 5.9 | 11.8 | 47.8 | 6.8 | 5.0 | 52.0 | 295.4 | 7.6 | 150.8 |
| Mean |  |  |  |  |  |  |  | 5862.3 | 11.2 | 17.1 | 3.3 | 5.3 | 69.9 | 0.2 | 1.4 | 0.8 | 2.6 | 0.2 | 0.7 | 0.2 | 0.5 | 0.3 | 2.1 | 4.1 | 0.5 | 0.2 | 6.8 | 65.8 | 4.6 | 27.0 |
| Standard deviation (n-1) |  |  |  |  |  |  |  | 3964.6 | 19.3 | 8.2 | 1.0 | 3.1 | 15.3 | 0.0 | 6.3 | 1.6 | 8.2 | 0.5 | 1.1 | 0.7 | 1.2 | 0.9 | 2.5 | 10.2 | 1.2 | 0.7 | 10.4 | 59.6 | 1.4 | 27.9 |

SI Table2: Information of each sample about sampling, HYSPLIT result, sequencing, meteorological data, source tracking results and alpha diversities
