## Supplemental Table3 for "Seasonal changes of airborne bacterial communities over Tokyo and influence of local meteorology"

| Taxonomy | type | Mann–Whitney U test |
| --- | --- | --- |
| Proteobacteria;Gammaproteobacteria;Enterobacteriales;Enterobacteriaceae;Citrobacter | non_continental | < 0.0001 |
| Actinobacteria;Actinobacteria;Corynebacteriales;Nocardiaceae;Gordonia | non_continental | < 0.0001 |
| Verrucomicrobia;Opitutae;Opitutales;Opitutaceae;Opitutus | non_continental | 0.012 |
| Proteobacteria;Betaproteobacteria;Burkholderiales;Burkholderiaceae;Burkholderia-Paraburkholderia | continental | 0.0002 |
| Firmicutes;Clostridia;Clostridiales;Clostridiaceae_1;Clostridium_sensu_stricto_1 | continental | 0.001 |
| Proteobacteria;Alphaproteobacteria;Rhodospirillales;Acetobacteraceae;Acidiphilium | continental | 0.005 |
| Bacteroidetes;Cytophagia;Cytophagales;Cytophagaceae;Spirosoma | continental | 0.003 |

SI Table 3: Results of Mann–Whitney U test for long transported genus
