## Supplemental Table4 for "Seasonal changes of airborne bacterial communities over Tokyo and influence of local meteorology"

| Taxonomy | Type | Mann-Whitney | BLAST relatives | Identity (%) | Source |
| --- | --- | --- | --- | --- | --- |
| ASV429: Parcubacteria;Candidatus_Adlerbacteria;NA;NA;NA | non-continental | 0.001 | HE650054 | 99.5 | MBR wastewater treatment pilot plant |
|  |  |  | JQ384174 | 97.1 | FACE soil sample |
| ASV609: Parcubacteria;NA;NA;NA;NA | non-continental | < 0.0001 | EU746773 | 99.8 | drinking water system |
|  |  |  | EF693410 | 96.8 | sediment |
| ASV1233: Parcubacteria;NA;NA;NA;NA | non-continental | < 0.0001 | LN528327 | 95.1 | drinking water |
|  |  |  | JQ781371 | 95.1 | stream sediment |
| ASV1036: Parcubacteria;NA;NA;NA;NA | non-continental | < 0.0001 | AB179672 | 98.5 | groundwater, 0.2 m-filterable fraction |
| ASV1556: Verrucomicrobia;Opitutae;Opitutales;Opitutaceae;Opitutus | non-continental | < 0.0001 | JQ655349 | 99.8 | drinking water |
|  |  |  | KX968744 | 99.3 | tropical urban freshwater |
| ASV137: Proteobacteria;Betaproteobacteria;Burkholderiales;Burkholderiaceae;Burkholderia-Paraburkholderia | continental | < 0.0001 | MG388565 | 100.0 | soil |
|  |  |  | KX509163 | 100.0 | rainwater |
|  |  |  | MG809290 | 100.0 | rice straw anaerobic digester |

SI Table 4: Result of Mann-Whitney U test and BLAST search for long transported ASVs
