## Supplemental Table5 for "Seasonal changes of airborne bacterial communities over Tokyo and influence of local meteorology"

|  | sample categories | all period | lower alpha cluster | higher alpha cluster |
| --- | --- | --- | --- | --- |
| Sample statistic (R): | HYSPLIT | 0.203 | 0.076 | 0.337 |
|  | Month | 0.331 | 0.192 | 0.396 |
| Significance level | HYSPLIT | 0.1 | 17.5 | 0.1 |
|  | Month | 0.1 | 6.3 | 0.1 |

SI Table 5: Results of ANOSIM between sample categories of HYSPLIT and Month
