## Supplemental Table6 for "Seasonal changes of airborne bacterial communities over Tokyo and influence of local meteorology"

| Meteorological factor | Pseudo-F | P(perm) |
| --- | --- | --- |
| Precipitation | 1.4819 | 0.023 |
| Air temperature | 1.6524 | 0.012 |
| **Wind speed** | **1.9559** | **0.001** |
| Radiation | 1.2255 | 0.114 |
| **Humidity** | **2.0737** | **0.003** |
| Wave_height | 0.74128 | 0.958 |

SI Table 6: Dissimilarities on the effects of meteorological factors by PERMANOVA. Bold shows significant dissimilarities.
